# The Central Nucleus of the Amygdala Encodes the Motivation to Pursue Ethanol

**DOI:** 10.64898/2026.05.11.724330

**Authors:** Matilde Castro, Jingren Gu, Teresa Dong, David J Ottenheimer, Celine Drieu, Patricia H Janak

## Abstract

Prior studies implicate the central nucleus of the amygdala (CeA) in reward motivation, yet how this region encodes motivation for ethanol (EtOH) seeking and consumption–and how this compares to encoding of natural rewards—remains poorly understood. We recorded single-unit neural activity from rats during a cued instrumental task in which lever insertion on each trial indicates the opportunity to lever press for ethanol and post-press lever retraction signals ethanol delivery. We found neurons responsive to multiple trial events (lever insertion and retraction cues, lever press, port entry, and reward licks), including neurons with rhythmic activity entrained to licking during EtOH consumption. Notably, CeA neural responses to the lever insertion cue at trial start encoded both the likelihood and speed of engagement during reward seeking. Using a supervised classifier, we found that trial engagement and motivation level could be decoded from pre-trial and lever insertion response periods. Finally, we assessed whether these effects generalized to natural reward seeking. Sucrose-rewarded rats (both ethanol-exposed and ethanol-naïve) showed higher motivation (more rewards earned and shorter first-press latencies) and stronger CeA recruitment and responses at lever insertion than ethanol-rewarded rats, in agreement with the notion that CeA circuits amplify reward-predictive cue signals to facilitate rapid action initiation under high motivational states. Together, these findings indicate that CeA neurons dynamically encode motivational states, with tonic activity and reward-predictive cue responses predicting both engagement in reward seeking and the vigor of action initiation. We suggest these patterns of neural activity drive ethanol and sucrose pursuit.

**Significance Statement:** How motivational states drive reward-seeking is a central question in systems neuroscience, yet the mechanisms linking neural signals to action initiation remain poorly understood. Recording single-unit activity in the amygdala central nucleus (CeA) during cued ethanol self-administration revealed that neural responses to reward-predictive cues encoded both the decision to engage in reward seeking and the vigor of action initiation, and motivational state was decoded from cue-evoked signals and tonic activity even before cue presentation. Consistent with greater motivation for the more palatable sucrose reward, CeA neurons showed stronger cue responses and greater anticipatory firing preceding tongue contact with reward, compared with ethanol-rewarded rats. These findings identify CeA activity patterns that link motivational state to the rapid initiation of goal-directed action.

## Introduction

Ethanol use is highly prevalent across many societies, with approximately 44% of people across the globe above the age of 15 reporting having consumed some ethanol in 2019 (World Health Organization, 2024). Preclinical studies of the underlying neurobiology of ethanol use have identified critical neural regions that influence ethanol self-administration, including the central nucleus of the amygdala (CeA). Initial studies in non-dependent rats found that CeA microinfusions of GABA_A_ and opioid receptor antagonists decrease ethanol self-administration (Heyser et al., 1999; Hyytiä & Koob, 1995), as does pharmacological inactivation and permanent lesions of the CeA (Möller et al., 1997; Roberts et al., 1996). Recent studies find CeA optogenetic activation and inhibition during ethanol consumption respectively increase and decrease ethanol intake (Fraser et al., 2024; Torruella-Suárez et al., 2020). While these findings collectively indicate that the CeA is involved in ethanol self-administration, the underlying neural activity patterns that support this behavior in real time have not been fully characterized.

The CeA is implicated in drug and natural reward seeking more broadly (Baxter & Murray, 2002; Cardinal et al., 2002; Gallagher & Holland, 1994; Janak & Tye, 2015; Warlow & Berridge, 2021). CeA optogenetic activation enhances seeking and intake of cocaine (Warlow et al., 2017), ethanol (Fraser et al., 2024) and sucrose (Fraser et al., 2024; Robinson et al., 2014) and CeA stimulation is itself reinforcing (Fraser et al., 2024; Torruella-Suárez et al., 2020). Neural recordings in rodents establish that CeA neurons respond to sucrose rewards as well as the cues that predict reward delivery (Muramoto et al., 1993; Shabel & Janak, 2009; Steinberg et al., 2020; Yang et al., 2023), in agreement with proposed roles of CeA in behavioral responding to motivationally-significant conditioned and unconditioned stimuli (Baxter & Murray, 2002; Cardinal et al., 2002; Gallagher & Holland, 1994; Janak & Tye, 2015; Warlow & Berridge, 2021).

We previously found that CeA activity was modulated during ethanol intake and that activity of CeA neurons is rhythmically aligned to licking of ethanol solutions (Fraser et al., 2024). Here, we built upon those findings and investigated CeA neuronal encoding of the motivation to pursue and consume ethanol and sucrose rewards. We recorded CeA single-unit activity during ethanol self-administration using a discrete-trials (DT) behavioral procedure in which lever insertion marked trial onset and the opportunity to lever press for ethanol, and lever retraction after lever presses signaled reward delivery; in the absence of responding, lever retraction marked trial termination. This design allowed us to compare CeA neural responses to lever insertion and retraction cues on trials in which subjects were, or were not, motivated to respond. Generalized linear models (GLM) identified CeA neurons significantly modulated to both cued (lever insertion and retraction) and self-generated (lever-press, port entry, licking) task events. We found that CeA neural responses to the lever insertion cue encoded whether rats choose to seek ethanol or abstain, and, on seek trials, how quickly they initiate ethanol seeking. Using a support vector machine (SVM), we found that seeking and motivation level could be decoded from neural activity in both pre-trial and lever insertion response periods. Finally, we assessed whether these effects generalized across reinforcement conditions. Sucrose-rewarded rats (both ethanol-exposed and ethanol-naïve) showed higher motivation and stronger CeA neural recruitment and responses at lever insertion than ethanol-rewarded rats, in agreement with the idea that CeA circuits amplify reward-predictive cue signals to facilitate rapid action initiation under high motivational states. Moreover, sucrose produced a shift in neuron modulation to licking, with a higher proportion of CeA neurons firing immediately before spout contact compared to ethanol, indicating stronger anticipatory coding under sucrose. Together, these findings show that CeA dynamically tracks within-in session changes in motivation and integrates predictive cues and consummatory events to drive ethanol and sucrose pursuit.

## Methods

We recorded neural data from three cohorts of rats. One cohort received pre-exposure to ethanol (EtOH) in the homecage and was rewarded with ethanol during training and recording sessions (EtOH/EtOH group). To provide a comparison between ethanol and natural reward neural correlates during the task, an additional cohort that was naïve to ethanol was rewarded with sucrose during training and recording sessions (NoEtOH/Suc). To control for possible effects of chronic ethanol on task neural correlates in general, a different cohort also underwent homecage pre-exposure to ethanol but was rewarded with sucrose (Suc) during training and recording sessions (EtOH/Suc). Methods were consistent across the three groups unless otherwise noted.

### Animal Subjects

Adult male and female (EtOH/EtOH: n = 12; 6F / 6M, EtOH/Suc: n = 4; 2F / 2M, NoEtOH/Suc: n = 6; 3F / 3M) Long-Evans rats (Envigo, Frederick, MD), weighing 200 to 275 g upon arrival, were single-housed in a temperature- and climate-controlled vivarium on a 12 h light:dark cycle. Food was available *ad libitum* except as noted during initial training (see Apparatus and Instrumental Training). Water was available *ad libitum*, and rats were provided with paper shredding enrichment in the homecage. Experimental procedures took place during the light phase and were performed in accordance with protocols approved by the Animal Care and Use Committee at Johns Hopkins University.

### Reward Solutions

#### EtOH/EtOH Cohort

Pure grain EtOH was diluted to 15% (v/v) in tap water for homecage exposure. For the behavioral task, 10% (v/v) EtOH was used as the reward.

#### EtOH/Suc and NoEtOH/Suc Cohorts

Sucrose (Fisher Bioreagents, Fair Lawn, NJ) was prepared as a 14.2% (w/v; isocaloric to 10% EtOH) solution in tap water.

## Homecage Ethanol Pre-exposure

To accustom rats to the taste and pharmacological properties of ethanol, rats from the EtOH/EtOH and EtOH/Suc cohorts were allowed to drink 15% EtOH freely in the home cage for 24 h on Monday, Wednesday, and Friday for 5 weeks. Rats had free access to water via a Lixit spout connected to the building water supply in the home cage. Ethanol bottles were weighed before and after each drinking session and rat weights were recorded before each drinking session to determine consumption in grams/kilogram. Rats advanced to instrumental training following consumption of 1 gram/kilogram of EtOH over the course of 24 h in at least 5 consecutive drinking sessions in the final week of exposure.

## Apparatus and Instrumental Training

Training and recording occurred in conditioning chambers housed within sound-attenuating boxes (Med Associates, St. Albans, VT) using MedPC (Med Associates, St. Albans, VT). Rats from the EtOH/EtOH group underwent a single 1 h port training session, in which 0.2 mL aliquots of 10% EtOH were delivered upon entry to the reward port situated in the center of the right chamber wall. Next, rats were trained to press a lever (Med Associates, St. Albans, VT) on the left side of the reward port for 5 daily ∼1 h sessions under continuous reinforcement (CRF) wherein one lever press was required for 0.2 mL 10% EtOH reward delivery. Sessions ended after rats had earned a maximum of 30 EtOH rewards or 1 h had elapsed. Rats were food restricted to 90% of their baseline weight for the first 3 of these CRF sessions to promote lever pressing and then allowed *ad libitum* food for remainder of training and recording sessions. Rats were next trained to perform the discrete trials task structure (DT1); in this task, trial onset was signaled by insertion of the lever. If rats successfully pressed the lever once following lever insertion, the lever was immediately retracted and ethanol was delivered in the reward port, and the trial was counted as a seek trial. Reward delivery was either synchronous with lever retraction (n = 6 / 12 EtOH/EtOH rats) or delivered 0.8 – 1.2 s after lever retraction to facilitate event separation in neural analyses (n = 6 / 12 EtOH/EtOH rats). Failure to complete the ratio within 60 s was considered an abstain trial; in these cases, the lever was retracted but no reward was delivered. The intertrial interval (ITI) was 30 s. After 5 DT1 sessions, the response requirement was increased to 3 lever presses (DT3) for an additional 5 sessions. In expert rats, DT3 sessions consisted of 50 trials.

EtOH/Suc and NoEtOH/Suc rats underwent the same training but were rewarded with 0.2 mL of 14.2% sucrose. For all sucrose rewarded sessions, there was a variable interval of 0.8 – 1.2 s between the lever retraction and reward delivery. As rats exhibited high motivation to respond for sucrose, sucrose DT3 sessions were lengthened to 100 trials to provide opportunity to detect within-session variations in motivation.

## Electrode Implantation

We constructed drivable electrode arrays consisting of a custom-designed 3D-printed drive and sixteen 50-µm insulated tungsten wires (A-M Systems), along with two silver ground wires (A-M Systems, Sequim, WA), soldered to two male 8-channel connectors (Omnetics Connector Corporation, Minneapolis, MN) that made up the headstage. For surgical implantation, rats were anesthetized with isoflurane (3 – 5% for induction, 1 – 2% for maintenance) and temperature was maintained at 37°C using a heating pad. Prior to surgery, rats were administered carprofen (5 mg/kg) and cefazolin (70 mg/kg). Arrays were advanced slowly (0.1 mm/min) to the left central amygdala (CeA; AP: −2.4 mm, ML: −4.2 mm, DV: −7.8 mm from bregma) via a 1 mm craniotomy window and secured to the skull with dental cement (Lang Dental, Wheeling, IL). Antibiotic and analgesic ointments were applied for infection and acute pain management, respectively. Rats recovered for at least 7 days following surgery before retraining sessions.

## Electrophysiological Recordings

For electrophysiological recordings, rats were tethered via a cable from their headstage to a commutator in the center of the chamber ceiling. After several retraining sessions to accustom rats to this tethering, recording sessions commenced. In the EtOH/EtOH group, recording sessions were about 62 min long, while in the EtOH/Suc and NoEtOH/Suc groups, sessions were 92 min long on average. During the session, continuous neural signals and task events time stamps were collected at 40kHz (OmniPlex system; Plexon, Dallas, TX) while rats performed the DT3 task, as in our prior studies (Cheng et al., 2025; Vandaele et al., 2019). Signals were high-pass filtered at 300 Hz, and spike events were detected using a voltage threshold set at 3 standard deviations above the mean of the continuous signal calculated over a 10 s pre-session period. For each session, we selected a channel with no discernable single-units to serve as a reference. After each recording, the electrodes were driven down by ∼100 μm and recording resumed in the new location at minimum 24 h later. Waveforms were manually sorted into units (Offline Sorter V4; Plexon, Dallas, TX) by using standard criteria based on waveform shape, stability, and refractory period violations (< 0.2% within 2 ms). Units not stably recorded throughout the full session were excluded. Behavioral event timestamps were parsed into task-relevant categories using NeuroExplorer 4.0 (Nex Technologies, Colorado Springs, CO) prior to analysis in MATLAB (MathWorks, Natick, MA). Only sessions with 10 or more reward deliveries were included for analysis.

For the EtOH/EtOH group, we obtained 851 neurons (71 ± 25 neurons, mean ± S.E.M., n = 12 rats) across 67 sessions. For analyses on the subset of neurons from EtOH/EtOH rats with corresponding video recordings, 457 neurons (76 ± 40 neurons, mean ± S.E.M., n = 6 rats) across 33 of the 67 sessions were obtained. For the EtOH/Suc group, we obtained 497 neurons (124 ± 62 neurons, mean ± S.E.M., n = 4 rats) across 29 sessions. For the NoEtOH/Suc group, we obtained 248 neurons (41 ± 16 neurons, mean ± S.E.M., n = 6 rats) across 30 sessions.

Baseline firing rates estimated from neuron ITI spike activity across the cohorts ranged from 0.07 to 44.3 Hz, with an average frequency of 3.9 ± 0.1 Hz.

At the conclusion of recordings, electrode sites were labeled by passing a DC current (50 μA, 30 s, cathodal) through all channels of each electrode prior to perfusion, and recording locations were verified to be in the CeA using previously described histological methods (Vandaele et al. 2019).

## Statistical Analysis

Summary statistics are represented as mean ± S.E.M. for Gaussian distributions (normality tested with a Shapiro-Wilk test) or median and interquartile range for non-Gaussian distributions (specified in the text). Paired data following a Gaussian distribution were compared using a paired t-tests. One-way ANOVAs were used for multi-group comparison of Gaussian distributions. Di□erences between groups were assessed using t-test with Bonferroni-correction for post-hoc analysis. Linear mixed-effects models (LMEs) were used to compare behavioral measures while accounting for repeated measures within subjects. Fixed effects were sex (when determining its effect on EtOH dosage or number of rewards acquired), or group and trial type (when determining their effect on movement variables), and random intercepts were included for each subject. Post-hoc comparisons were performed as planned contrasts derived from the linear mixed-effects models, with *p*-values corrected for multiple comparisons using the Bonferroni method. Derivations of behavioral data obtained from video recordings are described in Animal Position and Movement.

For comparisons of proportions of neurons encoding each event of interest relative to chance (5%), we used a two-sided exact binomial test. To compare multiple proportions across the 3 experimental groups, Chi-squared tests were used with Bonferroni corrections applied. All other neural encoding and decoding approaches as well as analyses of rhythmic neural activity are described in full detail below.

All statistical analyses were performed using MATLAB (MathWorks, Natick, MA). For electrophysiological data, analyses were performed on unsmoothed data. We did not explicitly calculate power or make use of statistical methods to predetermine sample size but based our sample size on our prior studies and previous publications. Data collection and analysis were not performed blind to the conditions of the experiments. All statistical tests were two-tailed.

## Animal Position and Movement

During electrophysiological recording sessions, behavior (all rats, except n = 6 EtOH/EtOH rats, 3F / 3M) was recorded at 60 frames/s using a Stingray camera (Allied Vision Technologies, Stadtroda, Germany). Video acquisition in CinePlex Studio (Plexon) was time-locked to neural recording using CinePlex Editor (Plexon, Dallas, TX). We used DeepLabCut (DLC; Mathis et al., 2018). to extract spatial coordinates of the headcap, the left and right ears of the animals, along with the middle tip of the lever when visible. We only used coordinates with a confidence level greater than 95%. Coordinate data were resampled to 1,000 Hz. Using these coordinates, we calculated instantaneous speed, distance from the lever, and angle of the head relative to the lever.

To calculate the instantaneous speed, we find the Euclidean distance between the headcap at *t* and *t* − 1 and divide it by Δ*t* (0.001 s).

To calculate the distance from the lever, we use the following formula:

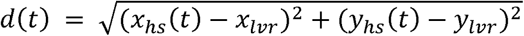

Where *x_lvr_* and *y_lvr_* are the coordinates of the lever and *x_hs_* and *y_hs_* are the coordinates of the headstage. Since the coordinate of the lever is stable across the session, *x_lvr_* and *y_lvr_* are defined as the average of all estimated *x* and *y* coordinates of the lever throughout the session.

To calculate the angle of the head relative to the lever, α_lever_, we used the area of the triangle formed by the ears and headcap to calculate an anchor point midway between the ears. With the anchor point, we construct two vectors: the first is from the anchor point to the headcap (the triangle’s height), while the second is from the anchor point to the lever. We use dot product to find the angle between these two vectors and use it as the angle of the head relative to the lever. The range of these angles is between 0° and 180°.

## Electrophysiology Data Analysis

### Seek Trial Encoding Model

To determine neural encoding of trial events and variables, we fit a kernel-based generalized linear model (GLM) to the spike counts of each neuron as in prior work (Fraser et al., 2025; Ottenheimer et al., 2023; Pillow et al., 2008; Vaccari et al., 2021). We first analyzed activity during rewarded (seek) trials.

## Data preparation

Spiking activity for each neuron *n* was binned into 25-ms windows, starting 1 s before lever insertion and extending 2 s after the final event of each trial, resulting in a spike count *y_n_(t)* for each neuron *n* at time *t*. The spike counts from seek trials were concatenated across trials to form a *T* × 1 vector for each neuron, where *T* represents the total number of time bins across seek trials. When modeling CeA spike counts during EtOH consumption, all available seek trials were used. When modeling CeA spike counts during sucrose consumption, the trials used were down-sampled within each session to match the average number of seek trials in ethanol sessions by randomly selecting 20 trials (∼average number of EtOH rewards) per session without replacement. This procedure was repeated across 100 iterations, with GLMs fit separately for each subsampled dataset, and variability across iterations retained for comparison analyses between experimental groups. Units with a firing rate below 1 Hz were excluded from GLM analysis (EtOH/EtOH: n = 43, EtOH/Suc: n = 18, NoEtOH/Suc: n = 17).

## Predictor matrix

The model included predictor kernels aligned to both externally controlled events (i.e. lever insertion, and lever retraction) and self-initiated actions (i.e. lever presses, rewarded port entries, port checks, individual licks, and lick bout initiations). Each event was associated with a fixed kernel window size (see Table 1.1), which defined the range of time lags relative to an event.

**Table 1.** Kernel window sizes for the GLM predictors. Time windows are expressed in ms relative to the onset of the corresponding event as time = 0.

| Event | Kernel Window (ms) |
| --- | --- |
| Lever Insertion | 0 – 500 |
| Lever Press (1 & 2) | -250 – 250 |
| Lever Retraction | 0 – 500 |
| Rewarded Port Entry | -500 – 500 |
| Unrewarded Port Entry | -500 – 500 |
| Individual Lick | -100 – 100 |
| First Lick in a Cluster | -100 – 100 |

We constructed a Toeplitz predictor matrix of size *T* × *l*, in which *T* is the total number of time bins and *l* is the number of lags required for the kernel. For each predictor, the predictor matrix contains diagonal stripes originating from the time an event occurs and 0 otherwise. The predictor matrices from seek trials were concatenated horizontally to yield a global prediction matrix *P* of size *T × L*, where *L* is the total number of predictor lags across events. Additionally, the model included trial-based covariates that identified the prior trial’s type (1 for seek, 0 for abstain) to capture trial-history effects, as well as the cumulative fraction of rewards to account for tonic changes in spike counts across the session. For neural recordings with corresponding video data, head angle relative to the lever and head speed were included as video-derived predictors to account for position- and movement-related modulation of neural activity, consistent with prior work incorporating kinematic variables into neural encoding models (Engelhard et al., 2019). Because these variables were strongly collinear, they were modeled as a single composite predictor. Prior to inclusion in the GLM, kinematic measures were resampled to match the temporal resolution of the neural data by binning to the same time bins used for GLM predictors and spike counts, as the original video-derived measures were sampled at a higher temporal resolution. Finally, spike history (for time bins *t* − 1 and *t* − 2) was included to model intrinsic temporal dependence in firing probability.

## Model Fitting

For each neuron *n*, the spike count *y_n_* at bin *t, y_n_(t)*, was modeled as:

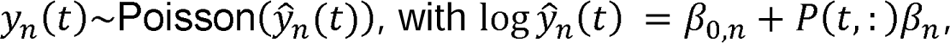

Where *β_0,n_* is the intercept representing the baseline log expected spike count per time bin of neuron *n, P(t,:)* is the global predictor matrix at time bin *t*, β_n_ is the vector for kernel weights for neuron *n* and ŷ_n_(t) is the model-predicted mean spike count from neuron *n* in bin *t*. The log link ensures positive predicted spike counts. The model was fit using the glmnet() package in MATLAB with ridge regularization (Friedman et al., 2010). The optimal regularization parameter λ was determined through 4-fold cross-validation for each by minimizing the negative log-likelihood on held-out data.

## Model Evaluation

Model performance was assessed by computing the variance explained using the formula:

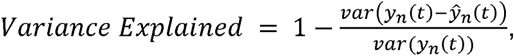

where *y_n_(t)* is the observed spike count and ŷ_n_(t) is the model-predicted spike count. To assess the contribution of each predictor to model variance, reduced models were fit by omitting event kernels, and the corresponding variance explained was computed.

To determine whether individual predictors significantly modulated neural activity, we used approaches described by (Fraser et al., 2025). For each neuron, we quantified the contribution of each predictor by computing the *F*-statistic comparing the full model to a reduced model in which that predictor was omitted:

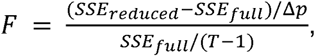

where *SSE* denotes the sum of squared errors, Δ*p* is the number of omitted predictors (i.e. number of predictors in full model – number of predictors in reduced model), and *T* is the number of time bins. Null *F*-statistics were obtained by fitting the same models to a null distribution generated by randomly resampling trial start times within the session boundaries while preserving the total number of bins per trial.

For each neuron, the observed *F*-statistics were evaluated against the null *F*-statistic distribution using Monte Carlo-derived p-values:

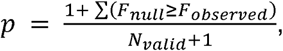

Where *N_valid_* denotes the number of valid null samples (variance explained greater than 0). For each neuron, a predictor was considered to significantly modulate activity if *p* < 0.05.

### Seek vs Abstain and Seek-fast vs Seek-slow Trial Encoding Models

To determine neural encoding of cue events based on trial type we fit a kernel-based GLM as in the Seek Trial Encoding Model, with the following modifications:

## Data preparation

Binned spike counts were constructed as in the Seek Trial Encoding Model, except that spike counts were concatenated across the 0.5 s following lever insertion and the 0.5 s following lever retraction. For the Seek vs Abstain Trial Encoding Model, concatenation was across all trials, whereas for the Seek-fast vs Seek-slow Trial Encoding Model it was restricted to seek trials. In contrast to the Seek Trial Encoding Model, no trial down-sampling was performed, and all available trials were used in this analysis.

## Predictor matrix

Predictor kernels and window sizes were constructed as in the Seek Trial Encoding Model, except that event timestamp data only included event occurrences binned in the 0.5 s following lever insertion and the 0.5 s following lever retraction. For the Seek vs Abstain Trial Encoding Model, concatenation was across performed across all trials, whereas for the Seek-fast vs Seek-slow Trial Encoding Model it was restricted to seek trials. Coincident events beyond lever insertion and lever retraction, together with their corresponding lags, were retained in the predictor matrix to account for explained variance but were not analyzed further. To model differences in cue-evoked activity across trial types, additional variables where lever insertion and retraction events were assigned a value of −1 or 1 depending on trial type (−1 = abstain, 1 = seek or −1 = seek-slow, 1 = seek-fast) were added. As in the Seek Trial Encoding Model, these event markers were expanded into lagged kernels via a Toeplitz construction, such that the trial type identity (−1 or 1) propagated across all lags of the corresponding event kernel. Thus, the resulting coefficients reflect trial type-specific modulation of cue-evoked activity across time. Trial-level covariates (prior trial type, current trial type, and trial number) were included to account for block-related changes in firing rate. Spike history for the *t* − 1 bin and *t* − 2 bin was included as in the Seek Trial Encoding Model.

## Model Fitting and Model Evaluation

For each neuron *n*, the spike count *y_n_(t)* was modeled, and the identification of significantly modulated neurons was evaluated as in the Seek Trial Encoding Model.

### Lick-modulation Analysis

We restricted lick-modulation analysis to licks emitted in lick trains, i.e. 2 or more licks with an inter-lick interval < 210 ms. Distributions of spike phases (in radians) were computed for each neuron (neurons with less than 50 spikes during the lick cycle were excluded: EtOH/EtOH, n = 21; EtOH/Suc, n = 42; NoEtOH/Suc, n = 15) and non-uniformity was tested using a Rayleigh test. Neurons with *p* < 0.01 were considered lick modulated.

### Classifier Analysis

We used a two-class support vector machine (SVM) to determine whether CeA neural activity could predict trial type, applied on spike counts in 25 ms time bins from −15 to +5 s from lever insertion. Spike counts were z-scored to the session-wide average by dividing each bin’s spike count by the mean spike count computed across the entire session binned activity. To avoid bias due to class imbalance, we performed stratified bootstrap resampling (n = 50; 9 seek and 9 abstain trials per session, or 9 seek-fast and 9 seek-slow per session). To establish significance relative to chance at the pointwise level, we computed Monte Carlo p-values (α = 0.05) at each time bin by comparing the mean decoding accuracy across resamples to a null distribution of mean decoding accuracy generated by shuffling trial type (n = 100). To assess significance in the 2.5 s before and after lever insertion, we computed the mean decoding accuracy across resamples within each window and compared it to a null distribution of the average decoder accuracy generated by shuffling trial labels (n = 100) in the pre- and post-lever insertion windows.

### Data Availability

The data and code used to analyze the data are available on GitHub: https://github.com/yerbamati/CeA_DT3_Modeling

## Results

### Rats exhibit robust but variable ethanol-seeking behavior in a discrete-trials procedure

Prior studies implicate the central nucleus of the amygdala (CeA) in ethanol seeking and consumption, yet how neural activity within this region encodes relevant events in real-time remains poorly understood. Here we obtained neural recordings from the CeA during ethanol self-administration using a behavioral procedure where subjects choose to obtain ethanol or not on each trial, thereby facilitating detection of motivation-sensitive neural activity. Rats were trained on a discrete trials (DT3) instrumental task in which lever pressing provided access to aliquots of ethanol. In each trial, a lever was inserted into the experimental chamber, and subjects were given 60 s to complete a fixed ratio of three presses (**Figure 1A**). Successful completion of the ratio resulted in lever retraction followed by ethanol delivery; these trials were classified as seek trials. Failure to complete the ratio within the allotted time resulted in lever retraction without reward delivery; these trials were classified as abstain trials.

**Figure 1.**
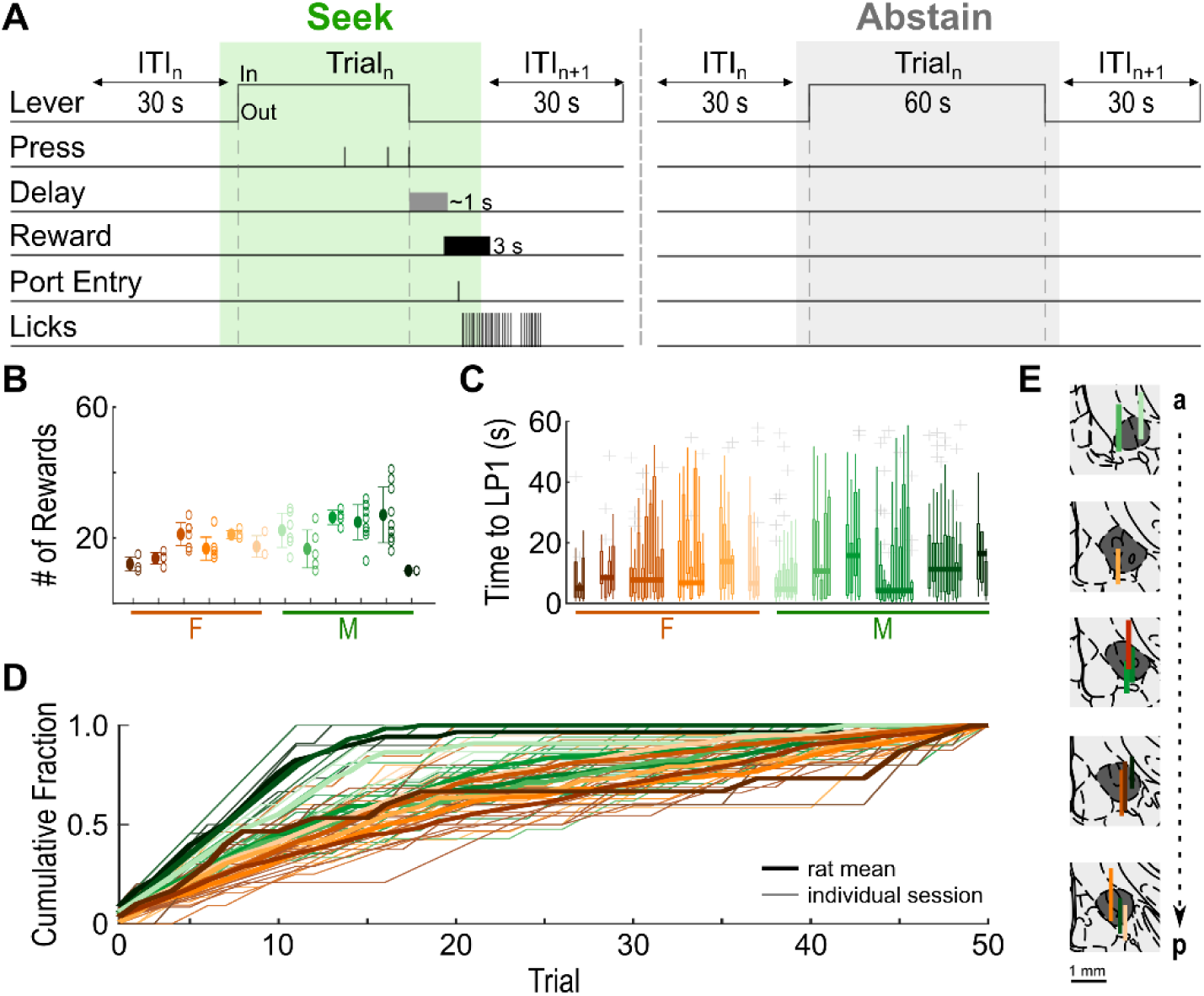
Rats exhibit robust but variable ethanol-seeking behavior in a discrete-trials procedure. **A.** Discrete Trials-3 (DT3) task design. Each trial is initiated by lever insertion after which the rat has 60 s to make 3 lever presses. If the rat completes the ratio, the lever is retracted, and 0.2 mL of liquid reward is delivered over 3 s after a variable time delay, and the trial is counted as a *seek* trial (marked with green). If the rat does not complete the ratio, the trial is counted as an *abstain* trial (marked with gray), the lever is retracted, and no reward is delivered. **B.** Number of EtOH rewards per session for each rat. Unfilled circles represent number of rewards attained by each rat during individual recording sessions. The across-sessions mean ± S.D. for each rat is denoted by a filled circle and error bars. **C.** Latency to first lever press across sessions for each rat. Rats are color-coded by individual and edges indicate 25^th^ and 75^th^ percentiles of the latency to press per recording session, with whiskers indicating most extreme datapoints that are not outliers. Outliers denoted with ‘+’ symbol. Thick horizontal lines indicate the median latency for each rat across all sessions. **D.** Cumulative fraction of rewards obtained by each rat per session (thin transparent lines) and by each rat on average (thick solid lines). **E.** Placements and drive path of electrode bundles for each rat. Scale bar = 1 mm.

During CeA recording sessions, rats earned on average 19.1 ± 1.6 rewards (mean ± S.E.M., n = 12 rats; **Figure 1B**), corresponding to 0.89 ± 0.07 g/kg ethanol consumed (mean ± S.E.M., n = 12 rats). Across sessions, ethanol intake did not differ across sexes (male, 6 rats: 0.79 ± 0.1 g/kg, n = 37 sessions; female, 6 rats: 0.98 ± 0.08 g/kg, n = 30 sessions; *t*(65) = −1.46, *p* = 0.15, linear mixed-effects model with subject as a random intercept).

When rats engaged in seeking, initial response latencies spanned a broad range within sessions (0.2 – 58.4 s; **Figure 1C**), indicating substantial variability in the initiation of goal-directed responding. The first lever press occurred on average 9.3 ± 1.2 s after lever insertion (mean ± S.E.M., n = 12 rats). Once initiated, responses were executed rapidly: with completion of the ratio in 1.4 ± 0.1 s (mean ± S.E.M., n = 12 rats), and reward port entry within 1.1 ± 0.1 s of lever retraction (mean ± S.E.M, n = 12 rats). Across sessions, rats responded on most trials early on, but cumulative responding gradually plateaued as session time progressed and animals increasingly abstained from responding (**Figure 1D**). This task design therefore enabled us to probe CeA activity across dynamic fluctuations in ethanol-seeking behavior.

### CeA neurons are recruited during ethanol self-administration

We recorded CeA single-unit activity during ethanol self-administration using drivable single-wire electrode bundles (**Figure 1E**). To assess the impact of behavioral events on CeA neural activity, we fit a kernel-based Poisson GLM to the spike train of each neuron binned in 25 ms windows (**Figure 2A**), restricting the analysis to seek trials. Predictors included task events (lever insertion and retraction, lever press, lick and port entry), and whole-trial variables, including the cumulative fraction of reward earned, to account for any drift in activity over the course of ethanol consumption, and prior trial type, to account for possible reward history effects. To capture short-term dependencies in firing, our model incorporated each neuron’s spike history over the preceding 50 ms (i.e. two time bins).

**Figure 2.**
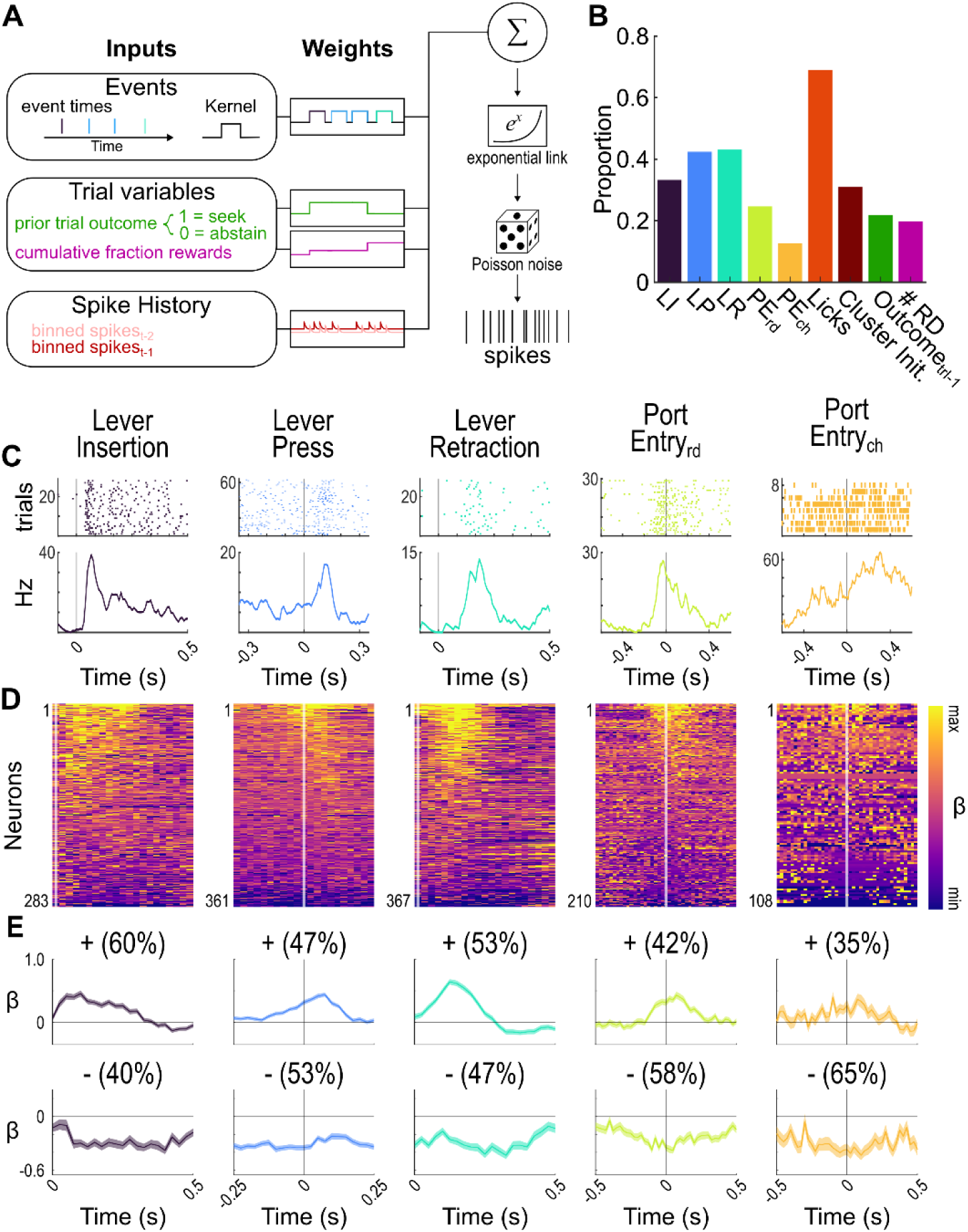
CeA neurons encode DT3 task events during seek trials. **A.** Schematic of encoding model quantifying the relationship between task variables and neural spike data. **B.** Proportion of neurons modulated by included predictors. LI, lever insertion; LP, lever press; LR, lever retraction; PE_rd_, rewarded port entry; PE_ch_, unrewarded port check; Cluster Init, first lick in a cluster; Outcome_trl-1,_ prior trial outcome (rewarded/unrewarded); #RD, number of rewards earned). **C.** Raster (top) and PSTH showing mean firing rate (bottom) for five events from example individual neurons determined to encode the event of interest. In raster plots, trials are sorted in ascending order starting at bottom of plot. **D.** Heat plots of β-weights estimated by GLM. Neurons are sorted along the y-axis by strength of modulation, with those with positive modulations (i.e. increases in spike count during an event kernel) at top and those with negative modulations (i.e. decreases in spike count during an event kernel) at bottom. The number of neurons encoding each task event is indicated at bottom of heat plots. Analysis interval time is indicated along x-axis in alignment with E. **E.** Mean β-weight for positively (top) and negatively (bottom) modulated neurons for each event of interest for analysis intervals depicted along x-axis, with event onset at t=0. The proportion of modulated neurons responding in each direction is noted. S.E.M. is indicated by a transparent shaded band.

Neurons in the CeA robustly encoded the main task events that occur during seek trials, with 699 neurons (82%) modulated by at least 1 event (**Figure 2B & 2C**). Examination of the β-weights, which reflect the degree of modulation in the kernel windows surrounding the events of interest, show a variety of event-related increases and decreases in neural activity, apparent across the population (**Figure 2D**) and in the population means (**Figure 2E**). Lever insertion, the cue signaling reward availability, drove changes in activity in 33% of the analyzed CeA neurons (**Figure 2B**), with 60% of those positively modulated (increased spike activity) and 40% negatively modulated (decreased spike activity) by this event (**Figure 2E**). An even greater proportion (43%) of neurons showed changes in activity related to lever retraction (**Figure 2B**), the event that signals imminent reward delivery. Significant modulations were also apparent across the range of emitted behaviors, including lever presses, port entries, and individual licks (**Figure 2B-E**). For all analyzed events, the proportions of modulated neurons were significantly higher than chance levels (lever insertion: k = 283, *p* = 1.0 × 10^-147^; lever press: k = 361, *p* = 1.3 × 10^-230^; lever retraction: k = 367, *p* = 1.5 × 10^-237^; port entry: k = 210, *p* = 7.8 × 10^-83^; port check: k = 108, *p* = 4.6 × 10^-18^; licks: k = 587, *p* ≈ 0, Binomial test, N = 851). Because many neurons encoded individual licks, we tested whether CeA activity was rhythmically aligned to licks during ethanol consumption. Rats licked at 7.8 ± 0.08 Hz (mean ± S.E.M., n = 12 rats; **Supplement 1A**), and 41% of neurons showed significant phase-locked firing by Rayleigh tests (**Supplement 1C & 1D**). The rhythmically aligned neurons showed a preferred phase centered near π/2, corresponding to activity just after tongue contact during retraction (**Supplement 1E-G**). When examining GLM whole-session predictors, we observed a small proportion of neurons (20%) that exhibited drift over the course of ethanol consumption, i.e. encoded the cumulative fraction of session rewards, and this proportion of modulated neurons was significantly higher than chance levels (k = 168, *p* = 5.8 × 10^-52^, Binomial test, N = 851). Furthermore, 22% of CeA neurons encoded recent task history, a proportion greater than chance (k = 185, *p* = 7.9 × 10^-64^, Binomial test, N = 851). These findings show that CeA neurons encode cue-, action-, consumption-, reward-, and recent history-related information, supporting a distributed representation of ethanol-seeking behavior.

We further explored how individual CeA neurons encode appetitive (lever insertion, lever press, and lever retraction) and consummatory (rewarded port entry and rhythmic licking) events. Neurons exhibited diverse response patterns to one or more events (**Figure 3A & B**), including activation, suppression, and mixed responses (**Figure 3C**). A majority (55%) of this population encoded two or more events (**Figure 3D**). About half (54%) of these neurons were modulated during both appetitive and consummatory phases, whereas fewer responded exclusively during the appetitive (30%) or consummatory (16%) periods (**Figure 3E**). Together, these results indicate that CeA neurons predominantly exhibit overlapping, multiplexed encoding across appetitive and consummatory events rather than forming discrete response classes.

**Figure 3.**
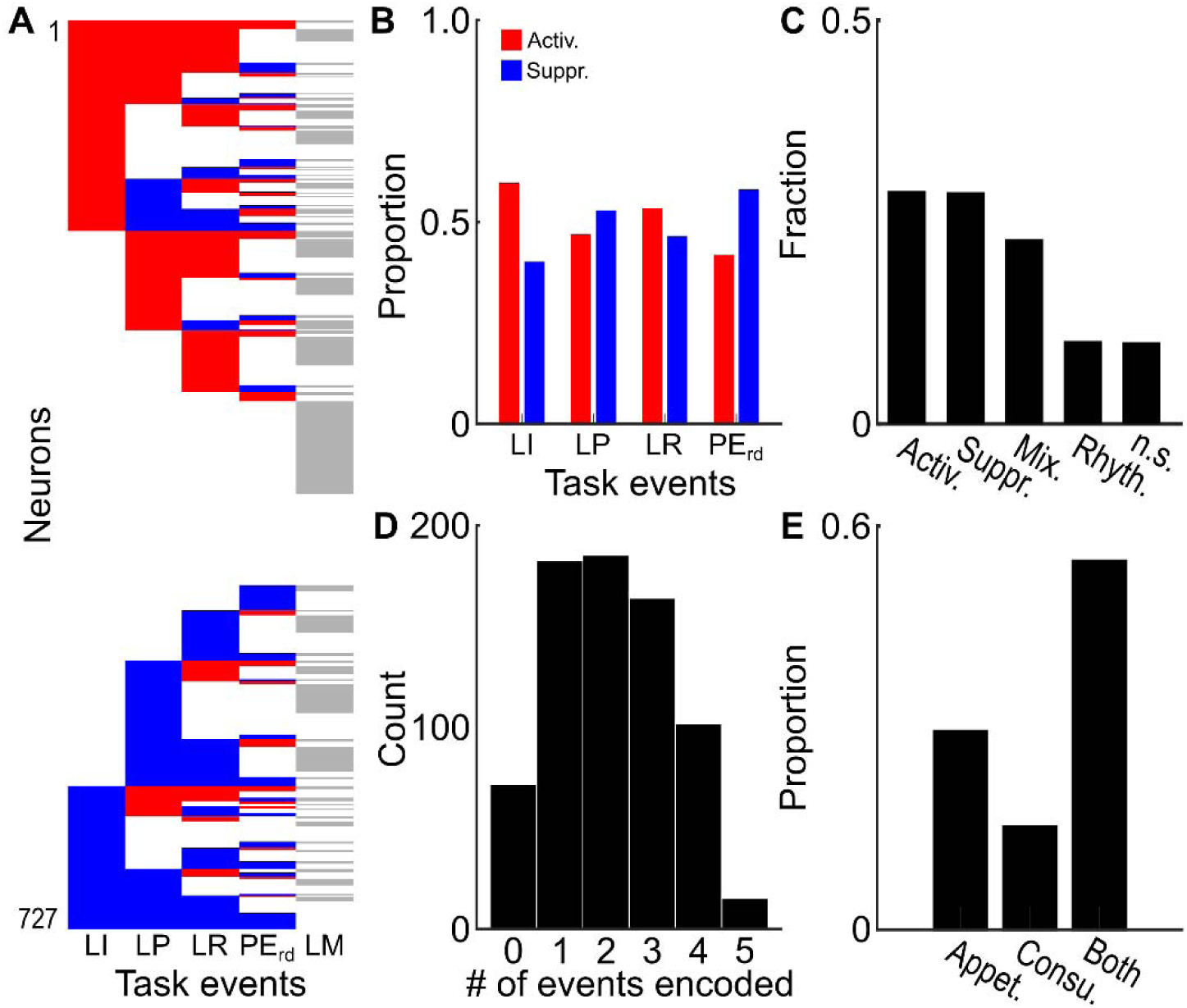
CeA neurons display heterogenous responses across events when rats seek EtOH. **A.** Responses of each neuron that underwent both GLM modeling and the Rayleigh test to events of interest (LI, lever insertion; LP, lever press; LR, lever retraction; PE_rd_, rewarded port entry; PE_ch_, unrewarded port check; LM, rhythmically lick-modulated)). Red = activation. Blue = suppression. Gray = lick modulated. Neurons sorted by first event encoded and direction of response, length of consecutive responses of same sign, and finally by presence of subsequent event responses. The purpose of this plot is to demonstrate the variety of multi-event encoding by individual units. **B.** Relative proportion of neurons that encode each event showing activation (red) or suppression (blue). **C.** Fraction of neurons that are only activated, only suppressed, only rhythmically modulated, show mixed responses, or do not significantly encode any event. **D.** Histogram of neurons by number of events each neuron encodes. **E.** Proportion of neurons that encode one or more events active during the appetitive phase, the consummatory phase, or both.

### Subsets of CeA neurons differentially encode reward-predictive cues according to motivational state

Because optogenetic activation of CeA during reward consumption rapidly alters motivated responding for a reward (Fraser et al., 2024; Robinson et al., 2014; Torruella-Suárez et al., 2020), we asked whether CeA neuronal responses vary based on real-time shifts in endogenous motivation. To address this question, we first assessed CeA responses to lever insertion and retraction cues, signaling reward availability and delivery, respectively, in seek versus abstain trials. A GLM (see Methods) revealed that 24% of CeA units responded to lever insertions regardless of trial type, 14% exhibited differential responses to lever insertion based on trial type, and 9% encoded lever insertion selectively for a single trial type (**Figure 4A**). These proportions were all above chance levels (lever insertion, general: k = 203, *p* = 2.5 × 10^-77^; lever insertion scaled by trial type: k = 116, *p* = 9.0 × 10^-22^; lever insertion only in certain trial types: k = 77, *p* = 1.2 × 10^-6^, Binomial test, N = 851). Thus, a subset of CeA neurons showed firing rate changes elicited by the lever insertion cue that varied depending on whether the rat chose to seek ethanol on that trial or not. In addition, most of the neurons encoding the lever retraction cue responded differently during seek or abstain trials (28%) (**Figure 4B**), with smaller subsets that responded to lever retraction overall regardless of trial type (14%), or only responded to lever retraction for one of the two trial types (14%). These proportions were also higher than chance (lever retraction, general: k = 116, *p* = 9.0 × 10^-22^; lever retraction scaled by trial type: k = 204, *p* = 6.9 × 10^-108^; lever retraction only in certain trial types: k = 123, *p* = 3.1 × 10^-25^, Binomial test, N = 851).

**Figure 4.**
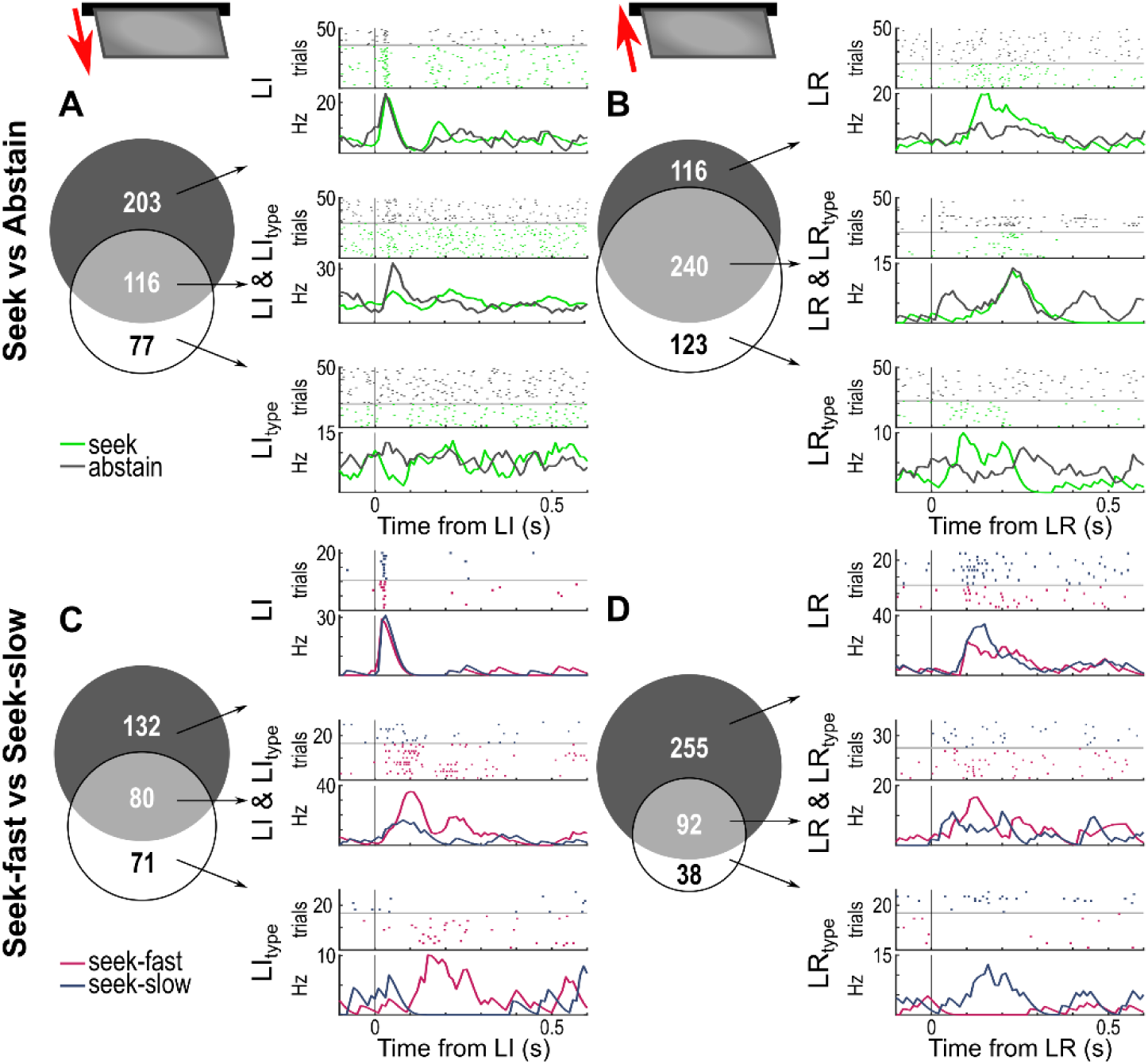
CeA neurons encode trial type and action vigor at cue events. **A.** Left, Venn diagram denoting number of neurons that encode the lever insertion cue generally (both seek and abstain trials; black circle), those that encode lever insertions differently for seek and abstain trials (gray overlap), and those that only encode trial type at ime of lever insertion (white circle) when the trial is a seek *or* abstain trial. Right, rasters (top) and PSTHs of mean firing rates (bottom) from example neurons of each type. Activity from seek trials shown in green, while activity from abstain trials shown in gray. In raster plots, trials are separated by trial type and then sorted in ascending order. **B.** As in **A**, but for lever retractions. **C.** As in **A**, but for seek-fast (denoted in magenta) or seek-slow (denoted in blue) trials. **D.** As in **B**, but for seek-fast or seek-slow trials.

The neural activity differences during seek and abstain trials indicated possible differences in cue encoding based on the level of motivation to seek the reward. To examine this possibility, we used each rat’s cross-session median latency to the first lever press to demarcate trials as seek-fast and seek-slow. Trials when a rat initiated the lever press sequence faster than the median latency were counted as seek-fast trials, and are taken to reflect relatively higher motivation, while those with a slower latency to first press than the median were counted as seek-slow trials and taken to reflect relatively lower motivation. Using the same logic as above for seek vs abstain trials, we applied a GLM (see Methods) to determine the effect of motivation to initiate a trial on lever-elicited neural activity. Of the recorded neurons, we found 16% encoded lever insertions generally, while 9% encoded lever insertion cues differentially for subsequent seek-fast versus seek-slow trials, and 8% significantly only encoded lever insertion on either seek-fast or seek-slow response trials (**Figure 4C & D**). These proportions were all above chance levels (lever insertion, general: k = 132, *p* = 5.4 × 10^-30^; lever insertion scaled by trial type: k = 80, *p* = 1.6 × 10^-7^; lever insertion only in certain trial types: k = 71, *p* = 5.0 × 10^-5^, Binomial test, N = 851). For lever retraction, 30% of neurons encoded this cue generally. Additional subpopulations encoded the level of motivation at the time of lever retraction, with 11% of the neurons encoding the event differently for seek-fast or seek-slow trials, and 4% encoding only seek-fast or seek-slow trials at the time of lever retraction. Numbers of neurons encoding lever retractions generally and scaled based on trial type exceeded chance levels, whereas the number of neurons encoding retractions selectively during seek-fast or seek-slow trials did not (lever retraction, general: k = 255, *p* = 2.8 × 10^-121^; lever retraction scaled by trial type: k = 92, *p* = 1.5 × 10^-11^; lever retraction only in certain trial types: k = 38, *p* = 0.53, Binomial test, N = 851). Overall, these findings support the proposal that cue-elicited spike activity in subpopulations of CeA neurons reflects real-time motivation to seek ethanol reward.

### CeA neural spiking predicts trial-by-trial performance

For half of the EtOH/EtOH rats (n = 6; 2F / 4M), we also obtained video recordings during neural data collection allowing us to quantify movement and head position across trial time. We focused our attention on these measures in the last 2.5 s of the intertrial interval (ITI) immediately before lever insertion, and the 2.5 s just after lever insertion, to understand how they might relate to motivation to seek ethanol in the different trial types.

Animals moved more during the ITI epoch preceding seek trials, and after lever insertion on seek trials, as reflected by higher mean ITI speed both before and after lever insertion compared to abstain trials (**Figure 5A**; Seek vs Abstain, pre lever insertion: *t*(5) = 4.3, p = 0.0079; post lever insertion: *t*(5) = 6.4, p = 0.0014, paired t-test). When considering the distance of the rat’s head relative to the lever at the pre- and post-insertion time points, no differences were observed for seek and abstain trials (**Figure 5B**; Seek vs Abstain, pre: *t*(5) = −0.27, *p* = 0.8; post: *t*(5) = −1.9, *p* = 0.11, paired t-test), while head angle relative to the lever was different after lever insertion (**Figure 5C**; Seek vs Abstain, pre: *t*(5) = −1.3, *p* = 0.25; post: *t*(5) = −4.8, *p* = 0.005, paired t-test).

**Figure 5.**
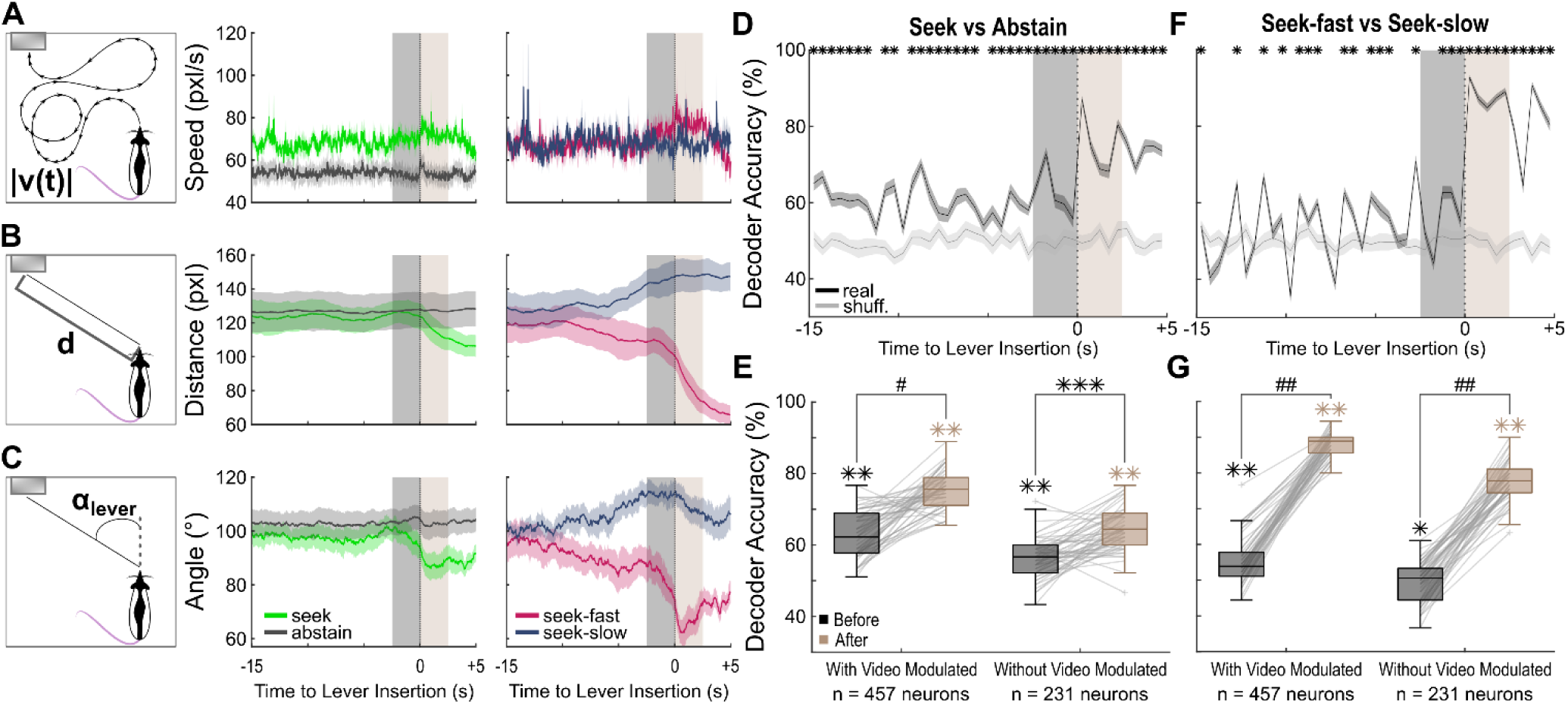
Pre-cue and cue-evoked CeA activity predict engagement and vigor of reward seeking independent of movement. **A.** Schematic (left) representing instantaneous speed, averaged by rat. Mean (± S.E.M.) instantaneous speed for −15 s to +5 s surrounding lever insertion shown for seek vs abstain trials (middle panel) and seek-fast vs seek-slow trials (right panel). Analysis windows for pre- and post-insertion comparisons indicated in gray and tan, respectively. **B.** As in **A**, but representing distance from the lever. **C.** As in **A**, but representing angle of the head relative to lever. **D.** Mean (± S.E.M.) accuracy across SVM resamples trained to separate neural data into seek and abstain trials across −15 s to +5 s surrounding lever insertion. Asterisks above time bins denote decoder accuracy significantly higher than shuffled decoder accuracy. **E.** Boxplots of decoder accuracy across resamples for seek vs. abstain trials in 2.5□s before (gray) and after (tan) lever insertion, comparing the full neuronal population with the subset excluding movement-modulated neurons. Resamples are connected by gray lines to show direction of effect. Box height indicates 25^th^ and 75^th^ percentiles, with whiskers indicating most extreme datapoints that are not outliers. Outliers denoted with ‘+’ symbol. Asterisks above boxes indicate significance vs. shuffled data, while asterisks above brackets denote significant differences between time windows for each group. * = *p* ≤ 0.05, ** = *p* ≤ 0.01, *** = *p* ≤ 0.001, # = *p* ≤ 1.0 × 10^-10^, ## = *p* ≤ 1.0 × 10^-20^. **F.** As in **D**, but the SVM is trained on seek-fast vs seek-slow trials. **G.** As in **F**, but using seek-fast vs seek-slow trial data.

In contrast, when focusing only on the seek trials and separating by response latency, ITI speed differed between seek-fast vs seek-slow trials only after lever insertion (**Figure 5A**; Seek-fast vs Seek-slow, pre-lever insertion: *t*(5) = 2.1, *p* = 0.088; post-lever insertion: *t*(5) = 3.0, *p* = 0.029, paired t-test). Examining head distance revealed differences at the pre-insertion time point and after lever insertion (**Figure 5B**; Seek-fast vs Seek-slow, pre: *t*(5) = −3.7, *p* = 0.014; post: *t*(5) = −6.9, *p* = 0.001, paired t-test). Similarly, differences in mean head angle relative to the lever between seek-fast and seek-slow trials were observed both before and after lever insertion (**Figure 5C**; Seek-fast vs Seek-slow, pre; *t*(5) = −7.3, *p* = 7.5 × 10^-4^; post: *t*(5) = −13.5, *p* = 4.0 × 10^-5^, paired t-test).

Overall, these behavioral findings show that speed differentiates seek vs abstain trials in the ITI, whereas measures of orientation/attention towards the lever distinguish different levels of motivation in trials in which rats choose to respond.

Because behavior differed across trial types in the ITI, and relative behavioral activation may reflect motivational state, we wondered if CeA neural activity during this period might also reflect motivation to respond for ethanol on the upcoming trial. To examine this issue, we asked whether CeA neural activity during the ITI, as well as following the lever insertion cue, could predict motivation to respond or abstain, and, for seek trials, to predict relative speed of responding. Focusing on the subset of EtOH/EtOH rats with corresponding video recordings, we applied a support vector machine (SVM) with 10-fold cross-validation trained on neural spike data that included 15 s of activity within the ITI prior to lever insertion and extended 5 s after lever insertion. Subsequent trial type—seek vs abstain—was decoded above chance levels at multiple time points within the ITI and, as expected, strongly after lever insertion. (**Figure 5D**). In contrast, the ability to decode if a subsequent trial would be seek-fast or seek-slow, was less stable during the ITI, sporadically exceeding chance levels, but increased greatly following the lever cue event (**Figure 5F**). Overall, the robust prediction following lever insertion indicates that population-level CeA activity at the time of the lever cue reflects ongoing motivational state, whereas significant but less stable decoding during the ITI suggests that baseline firing rates also carry motivational information about the upcoming trial prior to cue presentation.

We considered the possibility that trial motivation could be decoded during the ITI due to the presence of movement-correlated CeA neural activity. To determine how inclusion of neurons modulated by movement variables affect the performance of the decoder, we reran the SVM after removing the 44% of neurons determined to encode movement by a GLM performed on the subset of data with corresponding video recordings (**Supplement 2**) and again determined decoder accuracy. Comparisons were made between data in the 2.5 s before and after lever insertion for both the full population and the population excluding movement-modulated neurons and their relative null shuffled distributions. For the full population, decoder accuracy exceeded shuffle both before (*p* = 0.01) and after (*p* = 0.01; Monte Carlo method, 100 shuffles) lever insertion. When movement-modulated neurons were excluded, decoder accuracy remained significantly above shuffle before (*p* = 0.01) and after lever insertion (*p* = 0.01; Monte Carlo method, 100 shuffles). When comparing the average decoder accuracy in the full and reduced populations in the 2.5 s before and after lever insertion, we observed a significant effect of group, time relative to lever insertion, and an interaction between these two variables when decoding seek trials from abstain trials (Group: *F*(1,98) = 115.4, *p* = 3.0 × 10^-18^; Time: *F*(1,98) = 141.2, *p* = 1.1 × 10^-20^; Group x Time: *F*(1,98) = 7.7, *p* = 0.0067, ANOVA) reflecting the enhanced decoding when using the full population (**Figure 5E**). Overall, these findings show that CeA activity both before and after cue presentation can classify seek versus abstain trials on a trial-by-trial basis, even after accounting for movement-related modulation.

We found similar patterns in neural data from seek-slow or seek-fast trials (**Figure 5G**). For the full population, decoder accuracy exceeded shuffle before (*p* = 0.01) and after lever insertion (*p* = 0.01; Monte Carlo method, 100 shuffles). Similarly, for the population excluding movement-modulated neurons, decoder accuracy remained significantly above shuffle before (*p* = 0.02) and after lever insertion (*p* = 0.01; Monte Carlo method, 100 shuffles). Comparing the decoding accuracy of the full and partial populations, there were significant main effects of group and time relative to lever insertion, with a significant interaction between these variables (Group: *F*(1,98) = 118.6, *p* = 1.4 × 10^-18^; Time: *F*(1,98) = 1610.9, *p* = 1.2 × 10^-62^; Group x Time: *F*(1,98) = 9.5, *p* = 0.0026, ANOVA), again highlighting the enhanced decoding when the full population is considered. Together, these findings demonstrate that CeA population activity predicts both trial engagement and response vigor on a moment-to-moment basis and can do so independent of overt movement variables.

### Sucrose enhances motivation and cue-evoked CeA recruitment

We wondered whether the neural activity correlates observed in CeA during ethanol self-administration were similar when rats sought and consumed a natural reward. To test this, we recorded from two additional cohorts: rats with homecage ethanol exposure that received sucrose reward in the DT3 task (EtOH/Suc) and rats with no ethanol exposure that received sucrose reward (NoEtOH/Suc). The sucrose concentration we used is substantially more palatable than ethanol diluted in water and lacks ethanol’s pharmacological effects. Consistent with this, sucrose-rewarded rats performed more seek trials and very few abstain trials. To obtain enough of both trial types for neural analyses, we increased the number of trials per session. Behaviorally, sucrose-rewarded rats obtained more rewards than ethanol-rewarded rats by the 50th trial (*F*(2,19) = 77.5, *p* = 7.3 × 10^-10^) and initiated these seek trials more rapidly (*F*(2,19) = 23.8, *p* = 6.6 × 10^-6^, ANOVA) (**Figure 6A**), suggesting greater motivation for sucrose relative to ethanol. Overall reward consumption only differed by sex in the NoEtOH/Suc group (EtOH/Suc = male, 2 rats: 83.2 ± 6.8, n = 13 sessions; female, 2 rats: 73.1 ± 2.1, n = 16 sessions; *t*(27) = 1.65, *p* = 0.11; NoEtOH/Suc = male, 3 rats: 82.6 ± 6.1, n = 14 sessions; female, 3 rats: 66.2 ± 2.4, n = 16 sessions; *t*(28) = 2.9, *p* = 0.0067, linear mixed-effects model with subject as a random intercept). In contrast, other behavioral measures were similar across groups, including time to complete the ratio (*F*(2,19) = 1.6, *p* = 0.23), latency to enter the port (*F*(2,19) = 0.09, *p* = 0.92), and number of licks per reward (*F*(2,19) = 0.9, *p* = 0.42, ANOVA).

**Figure 6.**
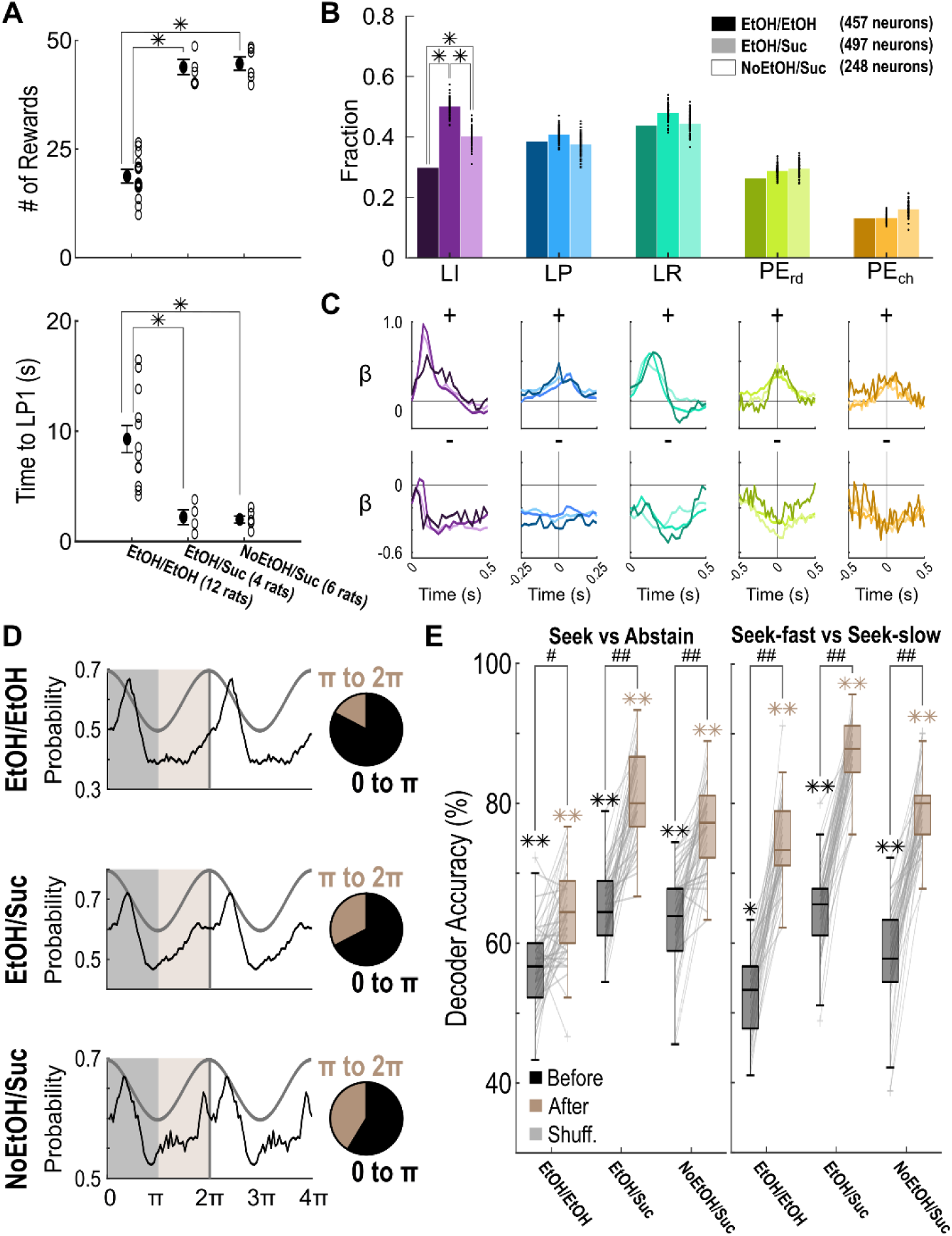
Higher motivation in sucrose-rewarded subjects is associated with enhanced CeA recruitment and anticipatory activity. **A.** Mean ± S.E.M. number of rewards and median latency to first lever press in 50 trials from the 3 groups. Bars with asterisks indicate significant differences, multiple comparisons test, Bonferroni correction applied. **B.** Fraction of neurons modulated by each event of interest (LI, lever insertion; LP, lever press; LR, lever retraction; PE_rd_, rewarded port entry; PE_ch_, port check). Fractions from each group represented using a gradient color scale. For the EtOH-rewarded group, the total fraction of neurons is represented by the darkest colored bar. Black dots on lighter-colored sucrose group bars denote the fraction of neurons encoding a task event for each of 100 subsampling iterations, while the bar denotes the mean fraction of neurons encoding an event. Asterisks indicate significant differences, multiple comparison tests, Bonferroni correction applied. **C.** Mean β-weights for neurons encoding each event. For EtOH-rewarded animals, the solid line represents mean across neurons encoding an event. For sucrose-rewarded animals, the solid line represents mean of 100 iterations. **D.** Mean spike probability of lick-modulated neurons across lick cycles (left) for EtOH/EtOH (top), EtOH/Suc (middle), NoEtOH/Suc (bottom) groups. Pie charts (right) depict proportion of neurons firing in 0 to π (black) or π to 2π (brown) phase of the lick cycle for each group. **E.** Boxplots of decoder accuracy across resamples for the three experimental groups in the 2.5□s before (gray) and after (tan) lever insertion when classifying seek vs. abstain trials (left) or seek-fast vs. seek-slow trials (right). Resamples are connected by gray lines to indicate direction of effect. Box heights indicate 25^th^ and 75^th^ percentiles, with whiskers indicating most extreme datapoints that are not outliers. Outliers denoted with ‘+’ symbol. Asterisks above brackets indicate significant differences between time windows for each group. * = *p* ≤ 0.05, ** = *p* ≤ 0.01, *** = *p* ≤ 0.001, # = *p* ≤ 1.0 × 10^-10^, ## = *p* ≤ 1.0 × 10^-20^. Neurons encoding video-derived variables were not included.

Investigation of the neural correlates revealed a pattern in agreement with this behavioral profile. A GLM analysis found largely similar event-related encoding patterns across reward types, with notable differences at trial initiation that may reflect the higher motivation observed in sucrose sessions (**Figure 6B**). Because sucrose sessions contained more rewards on average, we down-sampled sucrose trials to match ethanol sessions by randomly selecting 20 trials (∼average number of EtOH rewards) per session across 100 iterations. This allowed statistical comparisons of the proportions of event-modulated neurons across groups. The proportion of neurons modulated at lever insertion differed significantly across groups (*χ²*(2) = 40.9, *p* = 1.3 × 10^-9^, Chi-squared Test of Independence (3X2)) (**Figure 6B**), with each group’s proportion different from the other two. The EtOH/Suc group showed the largest fraction of lever insertion–modulated neurons, whereas the EtOH/EtOH group showed the smallest (EtOH/EtOH vs EtOH/Suc: *χ²*(1) = 40.9, *p* = 4.7 × 10^-10^; EtOH/EtOH vs NoEtOH/Suc: *χ²*(1) = 8.1, *p* = 0.014; EtOH/Suc vs NoEtOH/Suc: *χ²*(1) = 6.3, *p* = 0.035, Chi-squared Test of Independence (2X2), Bonferroni correction applied). In contrast, the proportions of neurons encoding other task events, namely lever presses, lever retractions, and port entries, did not differ across groups. Positively modulated neurons also exhibited larger cue-evoked responses in both sucrose-rewarded groups compared to the EtOH/EtOH group, whereas modulation strength for other task events was similar across reinforcers (**Figure 6C**).

CeA activity during licking was also phase-locked in sucrose sessions, broadly resembling the patterns observed during ethanol consumption. However, preferred firing phase differed across groups (*χ²*(2) = 35.4, *p* = 2.1 × 10^-8^, Chi-squared Test of Independence (3 x 2)), with a greater proportion of neurons in sucrose groups firing prior to tongue protrusion compared to the ethanol group (EtOH/EtOH vs EtOH/Suc: *χ²*(1) = 20.2, *p* = 2.1 × 10^-5^; EtOH/EtOH vs NoEtOH/Suc: *χ²*(1) = 31.7, *p* = 5.4 × 10^-8^). No significant differences were observed between the two sucrose groups (EtOH/Suc vs NoEtOH/Suc: *χ²*(1) = 3.3, *p* = 0.20, Chi-squared Test of Independence (2 x 2), Bonferroni correction applied), indicating that prior ethanol exposure did not alter phase locking during sucrose consumption (**Figure 6D**).

Together, these results indicate that CeA representations are largely conserved across reinforcers but are amplified at reward-predictive cues under higher motivational states.

### CeA population activity predicts trial outcome across reinforcement conditions

We wondered if trial engagement and response vigor could also be decoded on a per trial basis from CeA neural activity recorded in rats responding for sucrose. As with EtOH rats, rats responding for sucrose showed differences in movement and head position measures on trials with higher motivation, but there were notable differences between the EtOH- and sucrose-rewarded groups (**Supplement 3**). Given these motivation-related behavioral differences, we used a pointwise SVM decoder to predict trial type after excluding neurons modulated by speed and orientation across groups to reduce potential movement-related confounds. When decoding seek versus abstain trials, decoding was significantly better than shuffled for all groups both before (all *p*’s = 0.01) and after (all *p*’s = 0.01, Monte Carlo method, 100 shuffles) lever insertion. However, accuracy was higher in sucrose-rewarded rats than in ethanol-rewarded rats, with significant effects of group and time and a significant group x time interaction, arising from a larger improvement in decoding from the ITI to after the cue in sucrose rewarded subjects (Group: *F*(2,147) = 114.6, *p* = 1.0 × 10^-30^; Time: *F*(1,147) = 376.4, *p* = 2.3 × 10^-42^; Group x Time: *F*(2,147) = 14.2, p = 2.2 × 10^-6^, ANOVA) (**Figure 6F**, left). When decoding seek-fast versus seek-slow trials, significant decoding was detected relative to shuffled data both before (EtOH/EtOH. *p* = 0.02; EtOH/Suc & NoEtOH/Suc, *p* = 0.01) and after (all *p*’s = 0.01, Monte Carlo method, 100 shuffles) lever insertion. Again, decoder accuracy was higher for sucrose-rewarded rats, with significant main effects of group and time but no interaction (Group: *F*(2,147) = 112.1, *p* = 2.7 × 10^-30^; Time: *F*(1,147) = 1163.2, *p* = 1.0 × 10^-71^; Group x Time: *F*(2,147) = 0.1, *p* = 0.90, ANOVA) (**Figure 6F**, right). Thus, across reinforcement conditions, CeA population activity reliably predicts both trial engagement and response vigor, with stronger decoding performance in sucrose-rewarded rats.

## Discussion

Our experiments assessed CeA encoding of ethanol self-administration and differences in CeA neural activity based on current motivation to pursue reward. We found neurons encode both cue events and emitted behaviors when rats self-administer ethanol. Additionally, during ethanol consumption, some CeA neurons exhibited spiking patterns aligned with rhythmic licking. Neural activity elicited by lever insertion showed trial-by-trial variation accounted for by the real-time levels of motivation to lever press for ethanol. Across the population, CeA neural activity at the time of lever insertion significantly predicted whether rats would seek ethanol and, if so, how quickly, even when accounting for neural responses associated with movement. These findings were generally recapitulated in rats self-administering sucrose, although a greater degree of lever insertion cue encoding was observed, potentially in line with the higher motivational value of sucrose over ethanol in nondependent rats. Overall, our data join a growing body of evidence that the CeA is critical in driving reward seeking and intake for both drug and natural rewards (Cardinal et al., 2002; Douglass et al., 2017; Han et al., 2017; Hardaway et al., 2019; Janak & Tye, 2015; Mahler & Berridge, 2009; Robinson et al., 2014; Steinberg et al., 2020; Tom et al., 2019; Warlow et al., 2017; Warlow & Berridge, 2021). Our findings reveal that CeA neural activity encodes moment-to-moment changes in motivation to pursue an available reward and suggest that the neural mechanisms underlying CeA’s role in driving reward seeking include amplification of cue-elicited neural activity in higher motivational states.

Typically, studies on the CeA and ethanol focus on effects of dependence, where CeA-mediated changes in neural transmission are proposed to regulate negative affective states driving consumption (Funk et al., 2006; Gilpin et al., 2015; Koob, 2021; Koob & Volkow, 2016; Roberto, Madamba, et al., 2004; Roberto, Schweitzer, et al., 2004). However, ample evidence shows the CeA also regulates ethanol self-administration in non-dependent subjects (Barak et al., 2013; Fraser et al., 2024; Heyser et al., 1999; Hyytiä & Koob, 1995; Torruella-Suárez et al., 2020). We suggest that the CeA mechanisms underlying nondependent drinking are ‘co-opted’ by dependence-induced neural alterations to drive ethanol self-administration under those conditions, although this remains to be tested.

We observed robust engagement of the CeA by appetitive (lever cues and presses) and consummatory (port entry and reward licks) phases of behavior emitted within the ethanol self-administration task, as well as the parallel sucrose rewarded task. This is not surprising, given prior reports of CeA neural activity in response to cues, instrumental responses, and consumption for different reinforcers including sucrose, food, and intracranial electrical stimulation (Calu et al., 2010; Douglass et al., 2017; Fermani et al., 2025; Hardaway et al., 2019; Muramoto et al., 1993; Shabel & Janak, 2009; Steinberg et al., 2020; Torruella-Suárez et al., 2020; Yang et al., 2023). The present study found many similarities in neural correlates during ethanol and sucrose self-administration, in agreement with prior optogenetic findings in which channelrhodopsin-mediated activation of the CeA increased ethanol and sucrose intake (Fraser et al., 2024; Torruella-Suarez et al., 2020). However, we observed greater CeA engagement by the lever insertion cue that signaled opportunity to lever press for sucrosel, as indicated by a larger proportion of cue-modulated neurons and stronger phasic encoding immediately after lever insertion. Notably, sucrose rats responded for reward on more trials and showed faster response latencies than ethanol-reinforced rats. Therefore, greater CeA engagement by lever insertion in sucrose rats might reflect the higher motivation for sucrose than ethanol under the conditions used here, in line with the idea that CeA circuits amplify reward-predictive cue signals to facilitate rapid action initiation under high motivational states (Mahler & Berridge, 2009; Warlow & Berridge, 2021; Yang et al., 2023). However, we did not examine concurrent responding for these rewards by the same rat within a session, precluding direct comparison. Future within-session choice procedures could address this.

We did directly compare CeA cue responses during sessions for the same reward under conditions of higher or lower motivation. Because we used a discrete trials design where lever insertion and retraction occur on every trial regardless of behavior, we could compare cue-evoked neural responses on trials in which rats were motivated to lever press (seek trials) or not (abstain trials). CeA subpopulations responded differentially to lever insertion and lever retraction reward-availability cues depending on whether subjects were motivated to respond on that trial. In addition, cue responses were different for fast vs slow response latencies on trials in which subjects did seek reward. Finally, we found that seek vs abstain decisions and fast vs slow response latencies on seek trials were well-predicted by CeA cue-evoked activity across the population. Together, these findings suggest that the trial-by-trial real-time motivation to respond, as indicated by response likelihood and response vigor, is encoded by cue-elicited activity.

Neural activity during the ITI before lever insertion also reflected likelihood and vigor of future responding on that trial, although decoding was weaker than for cue-evoked activity. When motivation is relatively high, rats move more during the ITI, possibly engaging CeA movement-sensitive neurons. To discard the possibility that decoder accuracy in the ITI was driven by movement-correlated CeA neural activity, we decoded response likelihood or vigor after excluding movement-sensitive neurons and found ITI neural activity still predicted motivational state better than chance.

Blood and brain ethanol levels slowly accumulate during oral self-administration (Doherty & Gonzales, 2015; Doyon et al., 2005). To assess whether progressive ethanol exposure might influence CeA activity across the session, we included the cumulative fraction of ethanol rewards as a predictor in our GLM. A small proportion of recorded neurons displayed activity that varied with cumulative ethanol intake. Similar proportions were found in sucrose-reinforced groups, so it is unclear to what degree these session-wide changes in ethanol-reinforced sessions reflect pharmacological properties of ethanol. Instead, these drifts in firing rates may reflect other within-session factors, such as satiety or fatigue.

CeA neurons exhibited rhythmic modulation of spike activity in relation to licking for both rewards, matching our prior findings (Fraser et al., 2024). Lick-aligned neural activity is observed throughout the reward circuit, including within amygdala (Gutierrez et al., 2010), although presence specifically in the CeA has only recently been confirmed (Fraser et al., 2024). Peak rhythmic firing tended to occur as the tongue retracts into the mouth after liquid contact, presumably reflecting a time of sensation and perhaps outcome evaluation. While this prominent peak was observed for ethanol and sucrose reward, sucrose produced a shift in neuron modulation to licking, with a notable proportion of CeA neurons firing immediately before spout contact, suggesting stronger anticipatory coding under sucrose. Our finding of lick-entrained neural activity joins a growing body of work investigating CeA neural control of licking and biting of food and prey (Fermani et al., 2025; Han et al., 2017). Of note, we find individual neurons that respond to both appetitive (i.e., seeking) and consummatory aspects of behavior, highlighting the CeA as a region integrating both of these phases of behavior.

We included two groups of sucrose-trained rats, one receiving home-cage ethanol preexposure and one naïve to ethanol, to account for possible effects of chronic ethanol on CeA neural processing of reward seeking in general. Neural encoding in these two groups was similar, with some differences in encoding of the initial lever insertion cue. Specifically, while both sucrose groups had much higher proportions of neurons responding to the lever insertion cue at trial start compared to the ethanol group, the proportion of cue-responsive neurons in the sucrose group with home-cage ethanol preexposure was higher still than that of the ethanol-naïve group. Because of the small number of subjects in the ethanol preexposure group (n = 4), it is unclear if this distinction is meaningful. Of interest, we previously found effects of prior ethanol on cue encoding in the ventral pallidum, with a greater proportion of cue-evoked responses to sucrose cues that signal opportunity to seek reward after ethanol preexposure (Ottenheimer et al., 2019). In this past study as well as the current work, ethanol preexposure did not affect reward seeking behavior in the tasks used, yet evidence shows that chronic ethanol can alter neural activity and behavioral performance under conditions with greater cognitive challenge (Cheng et al., 2025). The possible cognitive/behavioral effect of modified neural processing of natural reward cues following chronic ethanol is an important area of continued investigation.

In sum, we found CeA neurons encode task-relevant stimuli and actions in rats self-administering ethanol or sucrose. Increased activity in response to cues that signal opportunity to respond for reward was observed for sucrose over ethanol and, for both rewards, the likelihood and vigor of reward-seeking behavior could be predicted trial-by-trial by neural responses to reward cues. Thus, CeA neural responses to reward stimuli reflect real-time variation in motivation, with an amplification of these signals to facilitate action initiation under high motivational states. Together, these findings show that CeA dynamically tracks within-session changes in motivation and integrates predictive cues and consummatory events to guide ethanol and sucrose pursuit.

## Acknowledgements

This research was supported by US National Institutes of Health grants R01AA027213 (PHJ), K99/R00DA059024 (CD) and F31AA030921 (MC). We thank all the members of the Janak laboratory for their input on this work.

## Author contributions

MC and PHJ designed research; CD and PHJ supervised the research; MC, JG, and TD performed research; DO provided GLM implementation guidance and preliminary code, refined for this data set by MC; MC, JG, and CD analyzed data; MC wrote original paper draft; MC, CD, and PHJ reviewed and edited the paper.

## Supplement for

**Supplemental Figure 1.**
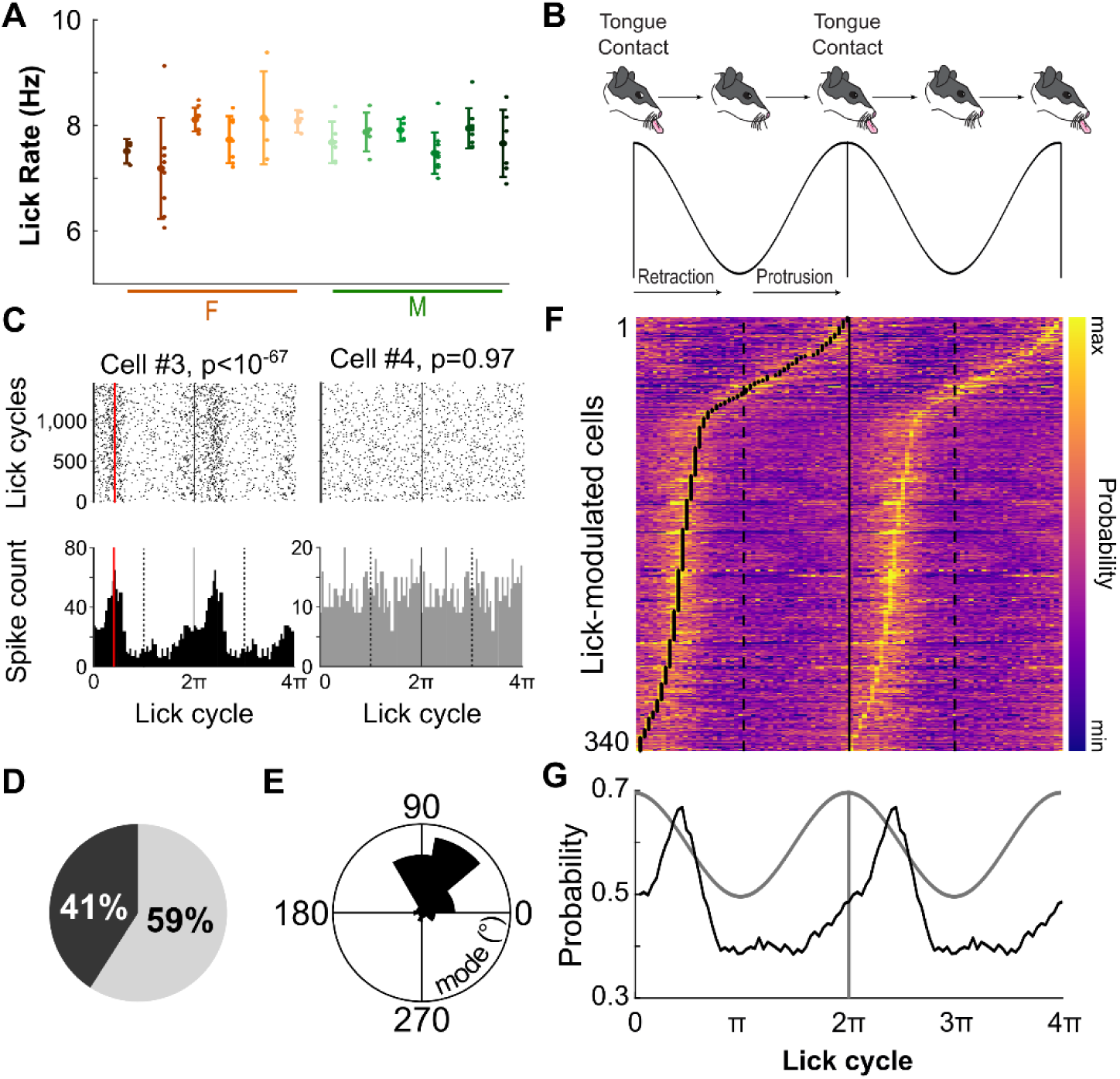
CeA neurons exhibit phase-locked activity to licking during EtOH consumption. **A.** Mean (± S.D.) lick rates for each subject during ethanol consumption for each DT3 recording session. Smaller symbols indicate lick rate for each individual session. **B.** Schematic representation of an individual lick cycle above its analogous sinusoidal rhythm. Times 0 and 2π indicate tongue contact with the liquid reward. **C.** Example spike rasters (top) and histograms (bottom) during lick cycles for two neurons recorded in the same session. The *p*-value of each neuron’s Rayleigh test is indicated. **D.** Proportion of neurons significantly modulated by licks (*p* < 0.01, Rayleigh test) indicated by black; unmodulated proportion indicated by gray. **E.** Circular histogram of the preferred firing phases. **F.** Heatmap of spike probability during lick cycles of all lick-modulated neurons. Black dots indicate the preferred firing phase (i.e., mode). **G.** Mean spike probability of lick-modulated neurons in **F** across lick cycles.

**Supplemental Figure 2.**
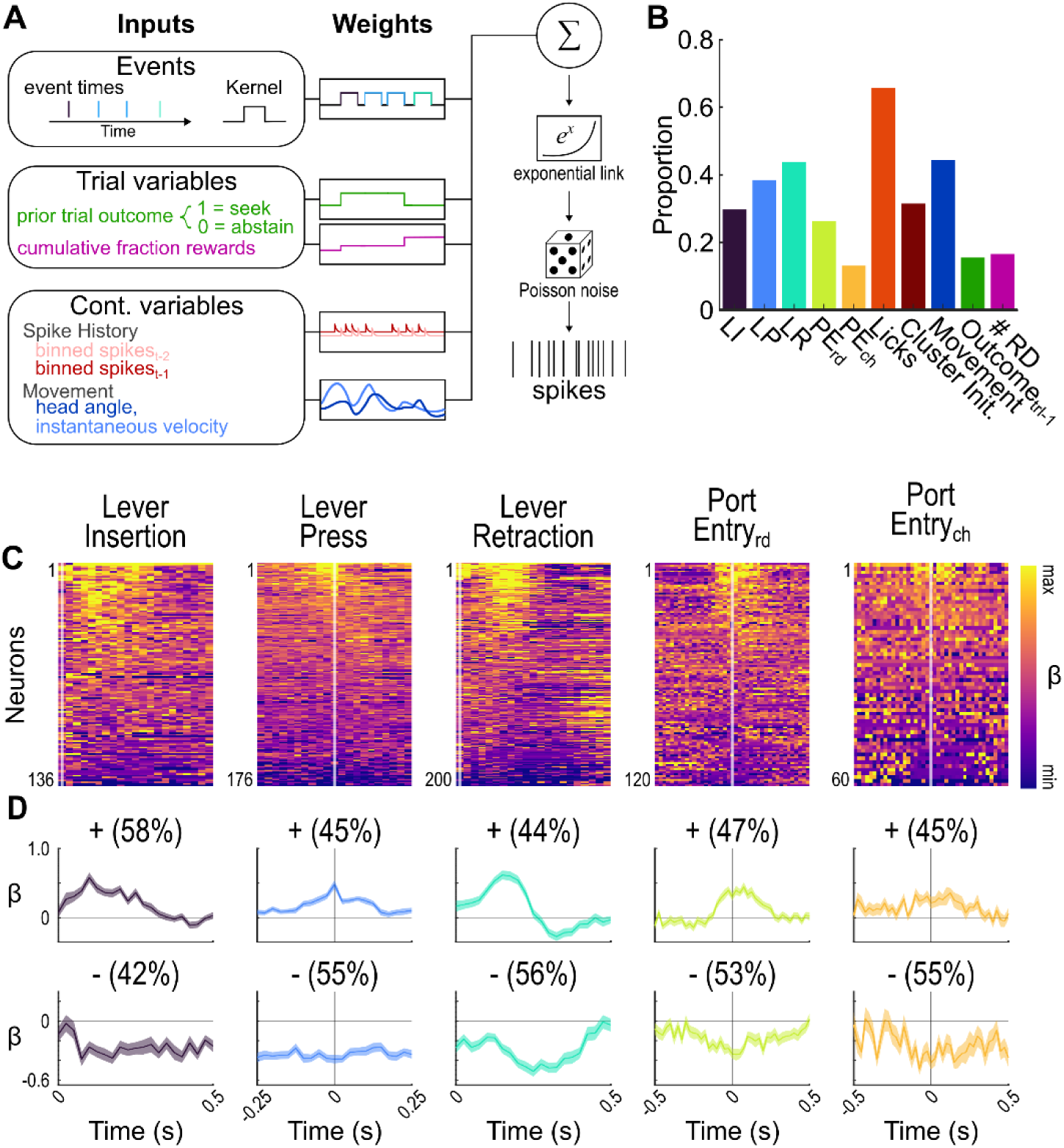
GLM results for subset of neurons with corresponding video data. **A.** Schematic of encoding model quantifying the relationship between task variables and neural spike data. **B.** Proportion of neurons modulated by the included predictors. **C.** Heat plots of β-weights estimated by the GLM. Neurons are sorted along the y-axis by strength of modulation, with those with positive modulations (i.e. increases in spike count during an event kernel) at the top and those with negative modulations (i.e. decreases in spike count during an event kernel) at the bottom. The number of neurons encoding each task event is reported at the bottom of the heat plots. The x-axis represents interval time as in D. **D.** Mean β-weights for positively (top) and negatively (bottom) modulated neurons for each event of interest. The proportion of modulated neurons responding in each direction is noted. The x-axis represents time, with event onset indicated by t=0. The S.E.M. is indicated by a transparent shaded band.

**Supplemental Figure 3.**
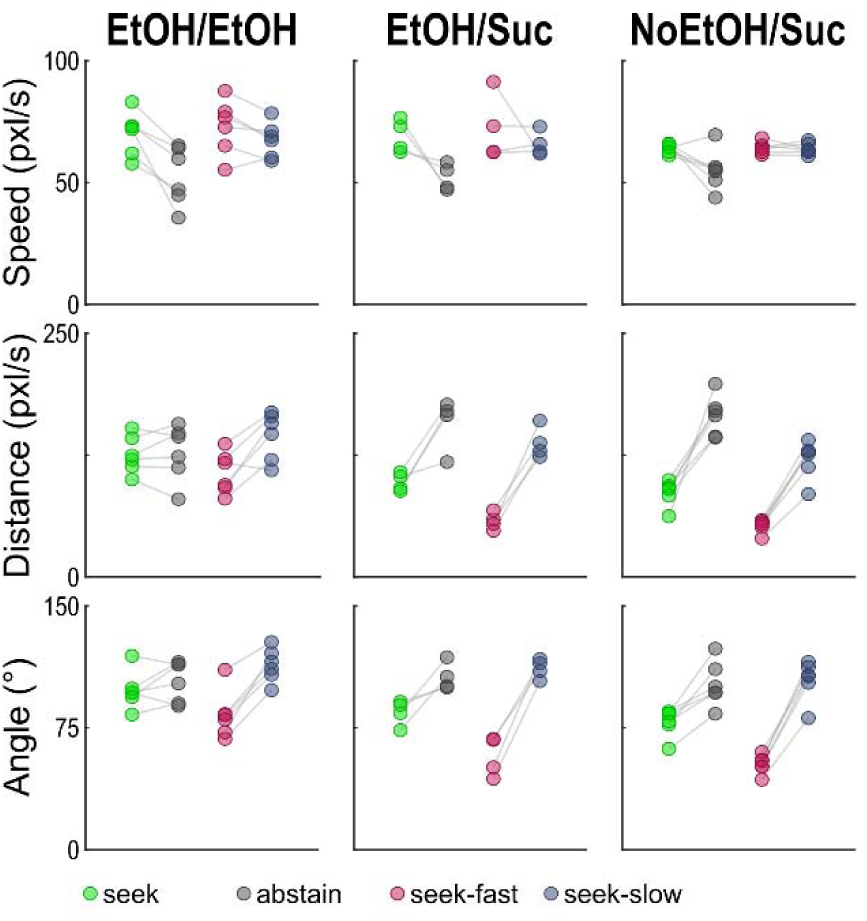
Behavioral differences prior to lever insertion reflect trial type, motivational state and action vigor. Behavioral variables (speed (top row), distance to lever (middle row), and head angle (bottom row)) in the pre-cue period for the three groups were analyzed using linear mixed-effects models (LME) with fixed effects of Trial Type and Group and random intercepts for subject. Omnibus fixed effects were assessed using F-tests from the mixed-effects model and are reported first, followed by Bonferroni-corrected planned simple effects contrasts to decompose significant interactions. These analyses found differences in movement variables in the ITI, as detailed below, suggesting that motivation to seek reward varied within the session, supporting the exploration of neural state encoding across trial types (Figure 6).

For movement speed, a linear mixed-effects model revealed a main effect of Trial Type (*F*(1, 7672) = 109.5, *p* = 1.9 × 10^-25^), no main effect of Group (*F*(2, 7672) = 0.13, *p* = 0.88), and a significant Trial Type x Group interaction (*F*(2, 7672) = 4.6, *p* = 0.01). Planned simple effects contrasts with Bonferroni correction showed a robust increases in speed during seek versus abstain trials for all three groups [EtOH/EtOH group (*t*(7672) = 10.5, *p* = 5.7 × 10^-25^), EtOH/Suc group (*t*(7672) = 9.6, *p* = 4.1 × 10^-21^), NoEtOH/Suc group (*t*(7672) = 7.3, p = 1.0 × 10^-12^), with lowest magnitude of effect in the NoEtOH/Suc group.

For head distance to lever, there was no overall main effect of Trial Type (*F*(1, 7672) = 0.002, *p* = 0.96), but a significant main effect of Group (*F*(2, 7672) = 7.8, *p* = 4.1 × 10^-4^), and a significant Trial Type x Group interaction (*F*(2, 7672) = 149.8, *p* = 1.5 × 10^-64^), indicating that the head distance in the two trial types differed across groups. Bonferroni-corrected planned simple effects contrasts found no effect of Trial Type in EtOH/EtOH (*t*(7672) = 0.05, p = 1.00), but significantly lower distance on seek trials in EtOH/Suc (*t*(7672) = −21.4, *p* = 2.8 × 10^-98^) and NoEtOH/Suc (*t*(7672) = −24.4, p = 1.1 × 10^-124^), indicating a trial type modulation of distance to the lever in sucrose-rewarded subjects.

For head angle relative to the lever, analysis revealed a significant main effect of Trial Type (*F*(1, 7672) = 5.5, *p* = 0.019), no main effect of Group (*F*(2, 7672) = 0.56, *p* = 0.57), and a significant Trial Type x Group interaction (*F*(2, 7672) = 13.4, *p* = 1.6 × 10^-6^). Bonferroni-corrected planned simple effects contrasts showed no difference in head angle based on trial type in EtOH/EtOH group after correction (*t*(7672) = −2.3, *p* = 0.057), but revealed significant effects in both sucrose groups (EtOH/Suc: *t*(7672) = −9.1, *p* = 2.6 × 10^-19^; NoEtOH/Suc: *t*(7672) = −9.8, *p* = 3.7 × 10^-22^), suggesting trial-dependent modulation of head orientation in the sucrose-rewarded subjects.

This analysis approach was also used to test for differences in movement variables between the fast and slow latency seek trials. For ITI speed, there was a main effect of Trial Type (*F*(1, 5216) = 9.3, *p* = 0.0023) and Group (*F*(2, 5216) = 3.8, *p* = 0.023), and a significant Trial Type x Group interaction (*F*(2, 5216) = 4.0, *p* = 0.019). Bonferroni-corrected contrasts revealed a significant difference in speed based on trial type for the EtOH/EtOH and EtOH/Suc groups (EtOH/EtOH: *t*(5216) = −3.05, *p* = 0.0067; EtOH/Suc: *t*(5216) = 3.7, *p* = 5.7 × 10^-4^), but no significant difference in speed based on trial type in the NoEtOH/Suc group (*t*(5216) = −0.5, *p* = 1.0). For head distance to lever, there was a significant main effect of Trial Type (*F*(1, 5216) = 88.6, *p* = 7.0 × 10^-21^) and Group (*F*(2, 5216) = 25.5, *p* = 9.5 × 10^-12^), and a significant Trial Type x Group interaction (*F*(2, 5216) = 36.7, *p* = 1.5 × 10^-16^). Post hoc comparisons revealed significant differences in distance from the lever for fast vs slows trials for all groups (EtOH/EtOH: *t*(5216) = 9.41, *p* = 2.1 × 10^-20^; EtOH/Suc: *t*(5216) = 32.6, *p* = 2.7 × 10^-211^; NoEtOH/Suc: *t*(5216) = 28.3, *p* = 9.9 × 10^-164^). For head angle, there was a significant main effect of Trial Type (*F*(1, 5216) = 101.8, *p* = 1.0 × 10^-23^), Group (*F*(2, 5216) = 17.4, *p* = 3.0 × 10^-8^), and a Trial Type x Group interaction (*F*(2, 5216) = 23.5, *p* = 6.6 × 10^-11^). Post hoc comparisons revealed significant differences based on trial type for all groups (EtOH/EtOH: *t*(5216) = 10.1, *p* = 3.1 × 10^-23^; EtOH/Suc: *t*(5216) = 29.6, *p* = 6.4 × 10^-178^; NoEtOH/Suc: *t*(5216) = 29.4, *p* = 2.6 × 10^-175^).

