## Supplemental Figures for "The Central Nucleus of the Amygdala Encodes the Motivation to Pursue Ethanol"

Contents:

Supplemental Figures 1-3

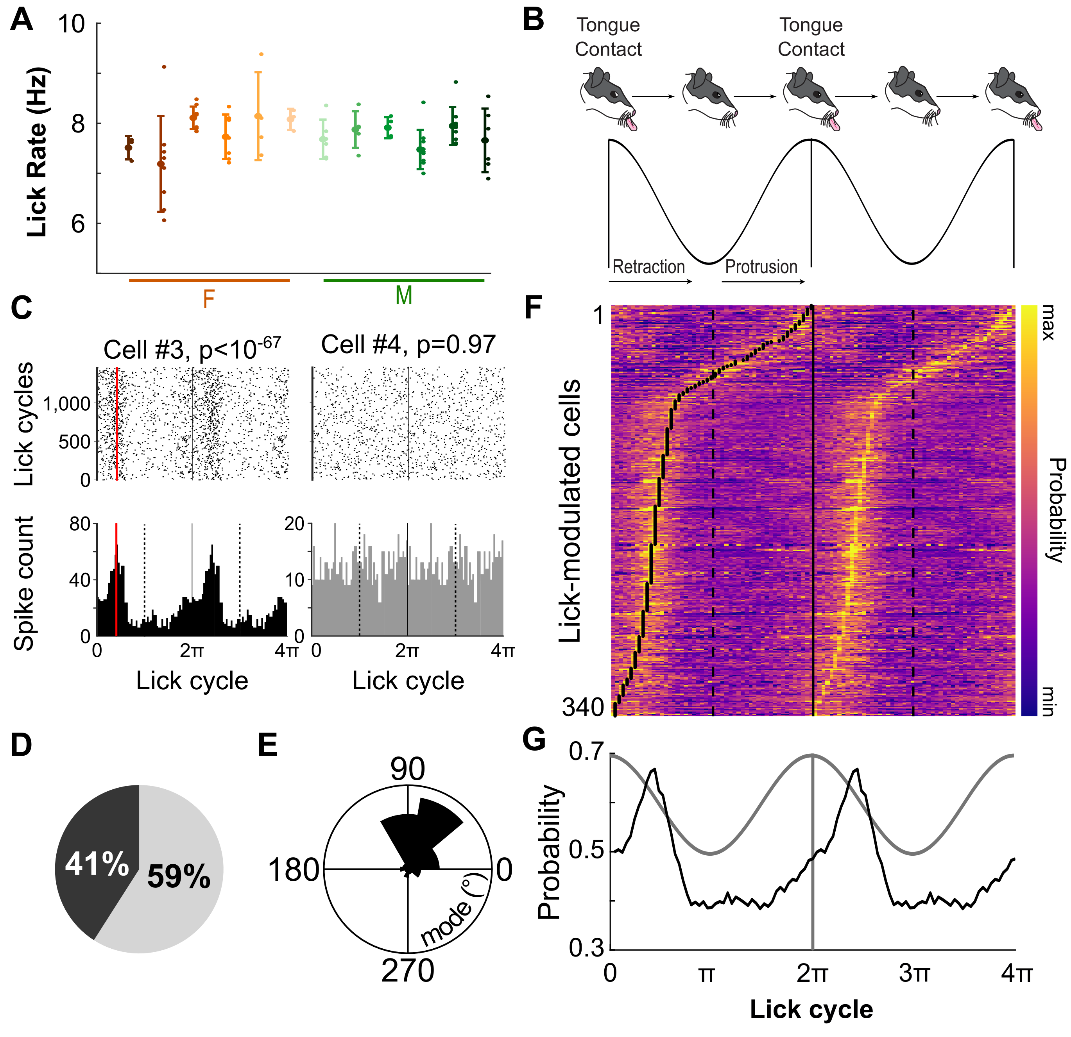

**Supplemental Figure 1.** **CeA neurons exhibit phase-locked activity to licking during EtOH consumption. A.** Mean (± S.D.) lick rates for each subject during ethanol consumption for each DT3 recording session. Smaller symbols indicate lick rate for each individual session. **B.** Schematic representation of an individual lick cycle above its analogous sinusoidal rhythm. Times 0 and 2π indicate tongue contact with the liquid reward. **C.** Example spike rasters (top) and histograms (bottom) during lick cycles for two neurons recorded in the same session. The *p*-value of each neuron’s Rayleigh test is indicated. **D.** Proportion of neurons significantly modulated by licks (*p* < 0.01, Rayleigh test) indicated by black; unmodulated proportion indicated by gray. **E.** Circular histogram of the preferred firing phases. **F.** Heatmap of spike probability during lick cycles of all lick-modulated neurons. Black dots indicate the preferred firing phase (i.e., mode). **G.** Mean spike probability of lick-modulated neurons in **F** across lick cycles.

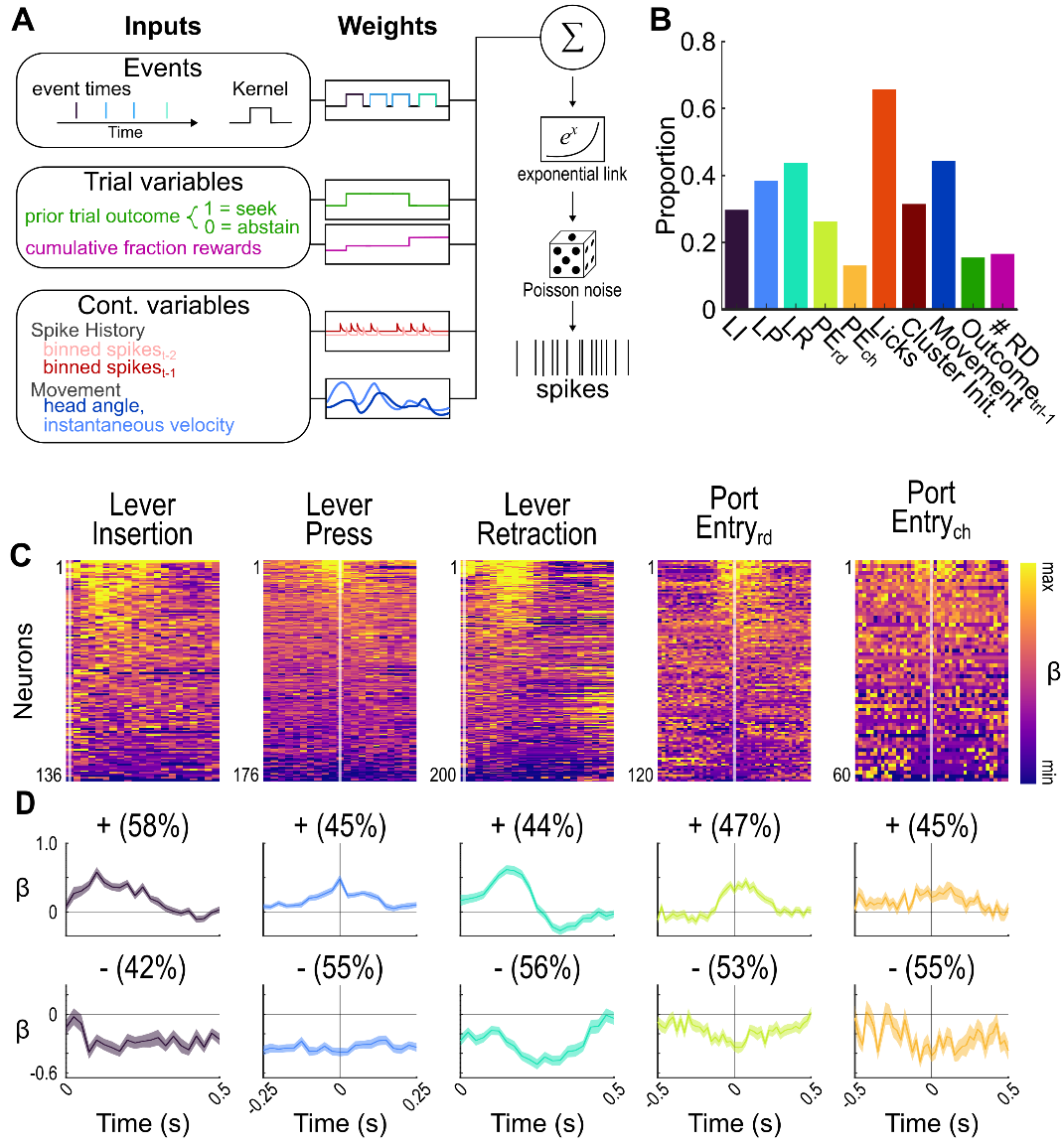

**Supplemental Figure 2.** **GLM results for subset of neurons with corresponding video data.** **A.** Schematic of encoding model quantifying the relationship between task variables and neural spike data. **B.** Proportion of neurons modulated by the included predictors. **C.** Heat plots of β-weights estimated by the GLM. Neurons are sorted along the y-axis by strength of modulation, with those with positive modulations (i.e. increases in spike count during an event kernel) at the top and those with negative modulations (i.e. decreases in spike count during an event kernel) at the bottom. The number of neurons encoding each task event is reported at the bottom of the heat plots. The x-axis represents interval time as in D. **D.** Mean β-weights for positively (top) and negatively (bottom) modulated neurons for each event of interest. The proportion of modulated neurons responding in each direction is noted. The x-axis represents time, with event onset indicated by t=0. The S.E.M. is indicated by a transparent shaded band.

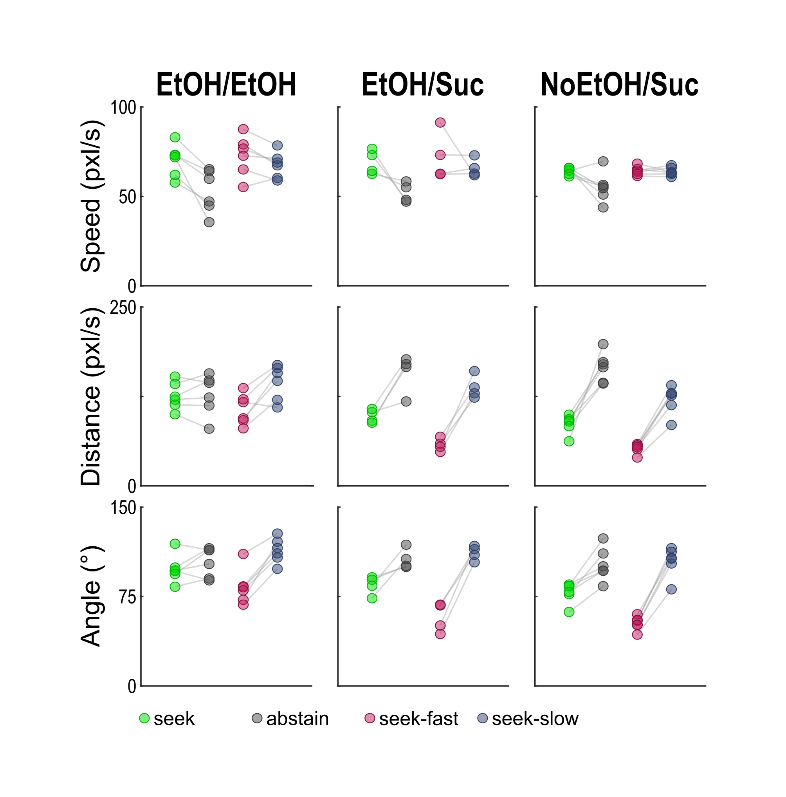

**Supplemental Figure 3.** **Behavioral differences prior to lever insertion reflect trial type, motivational state and action vigor.** Behavioral variables (speed (top row), distance to lever (middle row), and head angle (bottom row)) in the pre-cue period for the three groups were analyzed using linear mixed-effects models (LME) with fixed effects of Trial Type and Group and random intercepts for subject. Omnibus fixed effects were assessed using F-tests from the mixed-effects model and are reported first, followed by Bonferroni-corrected planned simple effects contrasts to decompose significant interactions. These analyses found differences in movement variables in the ITI, as detailed below, suggesting that motivation to seek reward varied within the session, supporting the exploration of neural state encoding across trial types (Figure 6).

For movement speed, a linear mixed-effects model revealed a main effect of Trial Type (*F*(1, 7672) = 109.5, *p* = 1.9 x 10^-25^), no main effect of Group (*F*(2, 7672) = 0.13, *p* = 0.88), and a significant Trial Type x Group interaction (*F*(2, 7672) = 4.6, *p* = 0.01). Planned simple effects contrasts with Bonferroni correction showed a robust increases in speed during seek versus abstain trials for all three groups [EtOH/EtOH group (*t*(7672) = 10.5, *p* = 5.7 x 10^-25^), EtOH/Suc group (*t*(7672) = 9.6, *p* = 4.1 x 10^-21^), NoEtOH/Suc group (*t*(7672) = 7.3, p = 1.0 x 10^-12^), with lowest magnitude of effect in the NoEtOH/Suc group.

For head distance to lever, there was no overall main effect of Trial Type (*F*(1, 7672) = 0.002, *p* = 0.96), but a significant main effect of Group (*F*(2, 7672) = 7.8, *p* = 4.1 x 10^-4^), and a significant Trial Type x Group interaction (*F*(2, 7672) = 149.8, *p* = 1.5 x 10^-64^), indicating that the head distance in the two trial types differed across groups. Bonferroni-corrected planned simple effects contrasts found no effect of Trial Type in EtOH/EtOH (*t*(7672) = 0.05, p = 1.00), but significantly lower distance on seek trials in EtOH/Suc (*t*(7672) = -21.4, *p* = 2.8 x 10^-98^) and NoEtOH/Suc (*t*(7672) = -24.4, p = 1.1 x 10^-124^), indicating a trial type modulation of distance to the lever in sucrose-rewarded subjects.

For head angle relative to the lever, analysis revealed a significant main effect of Trial Type (*F*(1, 7672) = 5.5, *p* = 0.019), no main effect of Group (*F*(2, 7672) = 0.56, *p* = 0.57), and a significant Trial Type x Group interaction (*F*(2, 7672) = 13.4, *p* = 1.6 x 10^-6^). Bonferroni-corrected planned simple effects contrasts showed no difference in head angle based on trial type in EtOH/EtOH group after correction (*t*(7672) = -2.3, *p* = 0.057), but revealed significant effects in both sucrose groups (EtOH/Suc: *t*(7672) = -9.1, *p* = 2.6 x 10^-19^; NoEtOH/Suc: *t*(7672) = -9.8, *p* = 3.7 x 10^-22^), suggesting trial-dependent modulation of head orientation in the sucrose-rewarded subjects.

This analysis approach was also used to test for differences in movement variables between the fast and slow latency seek trials. For ITI speed, there was a main effect of Trial Type (*F*(1, 5216) = 9.3, *p* = 0.0023) and Group (*F*(2, 5216) = 3.8, *p* = 0.023), and a significant Trial Type x Group interaction (*F*(2, 5216) = 4.0, *p* = 0.019). Bonferroni-corrected contrasts revealed a significant difference in speed based on trial type for the EtOH/EtOH and EtOH/Suc groups (EtOH/EtOH: *t*(5216) = -3.05, *p* = 0.0067; EtOH/Suc: *t*(5216) = 3.7, *p* = 5.7 x 10^-4^), but no significant difference in speed based on trial type in the NoEtOH/Suc group (*t*(5216) = -0.5, *p* = 1.0). For head distance to lever, there was a significant main effect of Trial Type (*F*(1, 5216) = 88.6, *p* = 7.0 x 10^-21^) and Group (*F*(2, 5216) = 25.5, *p* = 9.5 x 10^-12^), and a significant Trial Type x Group interaction (*F*(2, 5216) = 36.7, *p* = 1.5 x 10^-16^). Post hoc comparisons revealed significant differences in distance from the lever for fast vs slows trials for all groups (EtOH/EtOH: *t*(5216) = 9.41, *p* = 2.1 x 10^-20^; EtOH/Suc: *t*(5216) = 32.6, *p* = 2.7 x 10^-211^; NoEtOH/Suc: *t*(5216) = 28.3, *p* = 9.9 x 10^-164^). For head angle, there was a significant main effect of Trial Type (*F*(1, 5216) = 101.8, *p* = 1.0 x 10^-23^), Group (*F*(2, 5216) = 17.4, *p* = 3.0 x 10^-8^), and a Trial Type x Group interaction (*F*(2, 5216) = 23.5, *p* = 6.6 x 10^-11^). Post hoc comparisons revealed significant differences based on trial type for all groups (EtOH/EtOH: *t*(5216) = 10.1, *p* = 3.1 x 10^-23^; EtOH/Suc: *t*(5216) = 29.6, *p* = 6.4 x 10^-178^; NoEtOH/Suc: *t*(5216) = 29.4, *p* = 2.6 x 10^-175^).
